# *Phosphatidylinositol Transfer Protein-1* Integrates Insulin/IGF-1 and TOR Signaling to Negatively Regulate Lifespan and Healthspan in *Caenorhabditis elegans*

**DOI:** 10.1101/2025.11.28.691094

**Authors:** Yen-Hung Lin, Yun-Hsun Liao, Sin-Bo Liao, Tzu-Yu Lin, Muniesh Muthaiyan Shanmugam, Pei-Jia Hsu, Chang-Shi Chen, Tsiu-Ting Ching, Oliver Ingvar Wagner, Chiou-Hwa Yuh, Horng-Dar Wang

## Abstract

**Background:** *Phosphatidylinositol transfer protein-1* (*pitp-1*) is involved in phosphoinositide turnover. The role of *pitp-1* in promoting healthy longevity remains unknown. Our previous work showed that the phosphoinositide turnover genes *dagl-1* and *dgk-5* regulates lifespan, as overexpression of *dagl-1* or knockdown of *dgk-5* prolongs lifespan and enhances oxidative stress resistance through TOR signaling. As *pitp-1* is a key component of this pathway, we investigated its role in lifespan regulation and the underlying mechanisms, aiming to clarify whether it represents a critical regulator of healthy longevity and how it coordinates conserved signaling pathways to regulate aging.

**Methods:** *C. elegans* mutants, RNAi-mediated knockdown, and transgenic overexpression were applied to assess lifespan, motility, stress resistance. Temporal and tissue-specific RNAi were applied to identify the critical time window and tissue for *pitp-1*-mediated lifespan regulation. TOR signaling was measured by phosphorylated S6 kinase and puromycin incorporation, and transcriptomic analysis identified affected pathways.

**Results:** *pitp-1* negatively regulates lifespan and healthspan in *Caenorhabditis elegans*. Genetic deletion or RNAi-mediated knockdown of *pitp-1* extends lifespan, attenuates age-related motility decline, and increases oxidative stress resistance. Temporal and spatial analyses reveal that suppression of *pitp-1* in neurons during early adulthood is sufficient to promote healthy longevity. Mechanistically, these beneficial effects upon *pitp-1* reduction are mediated by suppressing TOR signaling. Conversely, *pitp-1* overexpression shortens lifespan and impairs healthspan via TOR activation. Moreover, *pitp-1* is transcriptionally repressed by DAF-16 downstream of insulin/IGF-1 signaling (IIS), and contributes to IIS-mediated lifespan extension.

**Conclusion:** These findings identify *pitp-1* as a novel regulator of healthy aging that integrates IIS and TOR pathways, providing new insights into conserved mechanisms for promoting healthy longevity.

## Introduction

Aging is an inevitable biological process with a progressive decline in physiological integrity, ultimately increasing vulnerability to stress and leading to death. It is a major risk factor for diverse aging-related diseases, including cancer, neurodegenerative disorders, metabolic syndromes, and substantially causes global healthcare burdens [1]. Thus, identifying genes and molecular mechanisms to promote healthy longevity is urgent for improving the quality of life in expanding aging populations.

Target of rapamycin (TOR) signaling and insulin/IGF-1 signaling (IIS) are evolutionarily conserved nutrient-sensing pathways in the regulation of aging [2, 3, 4]. TOR signaling integrates metabolic cues to control protein synthesis, growth, and metabolism [2, 5]. Suppression of TOR activity reduces phosphorylated S6 kinase (p-S6K) levels, lowers protein translation, dampens anabolic signaling, extends lifespan and enhance stress resistance across species [6, 7]. Similarly, reduced IIS signaling suppresses the PI3K–PDK–AKT kinase cascade and prevents the phosphorylation of DAF-16/FOXO by phosphorylated AKT (p-AKT), which allows DAF-16/FOXO to translocate into nucleus and orchestrate gene expression program that protects against cellular damage, improves stress resistance, maintains homeostasis and promotes healthy longevity. Notably, these two pathways are functionally interconnected. IIS-activated p-AKT inhibits the TSC1/2, which prevents Rheb from the repression by TSC1/2 and activates TOR complex 1 (TORC1) [3, 8]. Conversely, TOR complex 2 (TORC2) can phosphorylate and activate AKT [9]. This crosstalk fine-tunes cellular and physiological responses to metabolic conditions, highlighting a coordinated regulatory network that governs the aging process. Understanding the integration of TOR and IIS signaling offers crucial insights into the molecular logic of aging and provides promising targets for interventions aimed at extending lifespan and delaying age-associated physiological decline.

Phosphatidylinositol transfer proteins (PITPs) are conserved lipid transfer proteins that mediate the transport of phosphatidylinositol (PI) and phosphatidic acid (PA) between the endoplasmic reticulum (ER) and plasma membranes (PM), playing essential roles in phosphoinositide (PPI) turnover and maintaining PIP2 homeostasis [10, 11]. PITPs are conserved across species with homologs among *C. elegans*, *Drosophila,* and mammals [11, 12], and classified into two evolutionarily conserved classes of PITPs, class I and class II [12]. In *C. elegans*, *pitp-1*, the sole class II PITP, is mainly expressed in sensory neurons and regulates chemotaxis in response to environmental cues [13]. It also facilitates rapid recovery of feeding behavior after hypoxia by limiting diacylglycerol (DAG) availability and suppressing PKC activity in *mod-1*-expressing neurons [14]. In *Drosophila*, the class II PITP ortholog *rdgB* is essential for PPI cycling during phototransduction in photoreceptor cells [15]. In mammals, Nir2, homologous to *pitp-1*, binds PA and enhances the MAPK and PI3K/AKT signaling pathways in response to growth factor stimulation [16]. Suppression of Nir2 reduces breast cancer cell migration and metastasis [17]. Despite the roles of PITPs in neuronal signaling and cancer biology, whether *pitp-1* contributes to lifespan regulation remains unknown.

Our previous study demonstrated that overexpression of *dagl-1*/*inaE* (DAG lipase) or knockdown of *dgk-5*/*rdgA* (DAG kinase) extends lifespan and enhances oxidative stress resistance in *C. elegans* and *Drosophila*, likely through reduced levels of PA and subsequent inhibition of TOR signaling [6]. In addition, the phospholipase C β (PLCβ) homolog *egl-8*, another PPI cycle component, has been shown to regulate lifespan in *C. elegans*, as the null mutant *egl-8(n488)* exhibits extended longevity [18]. These findings suggest that components of the PPI cycle may play a role in lifespan regulation. Intriguingly, our previous multiple stress genetic screening by oxidative stress and starvation to find the mutants with increased stress tolerance as longevity candidates not only identified the mutant with *inaE* up-regulation in *Drosophila* [6] but also uncovered the mutant of *rdgB* (unpublished data), the ortholog of *pitp-1*. This prompts us to examine whether *pitp-1* participates in lifespan regulation and stress response in *C. elegans*. Here, we demonstrated the reduction of *pitp-1* promotes healthy longevity via modulating TOR signaling. Furthermore, *pitp-1* acts downstream of DAF-16 and contributes to IIS-mediated lifespan regulation, suggesting that *pitp-1* functions as a potential integrator of IIS and TOR signaling crosstalk. The transcriptomic analysis supports this notion, as *pitp-1* reduction is associated with gene expression changes that resemble those predicted for reduced TOR and IIS signaling activity. Together, our findings uncover *pitp-1* as a novel regulator of healthy longevity and provide new insights into conserved mechanisms of lifespan regulation.

## Methods

### C. elegans strains

*C. elegans* were maintained at 20°C on NGM agar plates seeded with *E. coli* OP50 using standard protocols. The following alleles and strains were used in this study: Wild-type Bristol N2, JN1297: *pitp-1(pe1297)*, *pitp-1(tm1500)*, TU3311: *uIs60 [unc-119p::YFP + unc-119p::sid-1]*, TU3401: *sid-1(pk3321); uIs69 [pCFJ90 (myo-2p::mCherry) + unc-119p::sid-1]*, WM118: *rde-1(ne300); neIs9 [myo-3::HA::RDE-1 + rol-6(su1006)]*, VP303: *rde-1(ne219); kbIs7 [nhx-2p::rde-1 + rol-6(su1006)]*, XE1375: *wpIs36 [unc-47p::mCherry] I. wpSi1 [unc-47p::rde-1::SL2::sid-1 + Cbr-unc-119(+)]; eri-1(mg366); rde-1(ne219); lin-15B(n744)*, XE1474: *wpSi6 [dat-1p::rde-1::SL2::sid-1 + Cbr-unc-119(+)]; eri-1(mg366); rde-1(ne219); lin-15B(n744)*, XE1581: *wpSi10 [unc-17p::rde-1::SL2::sid-1 + Cbr-unc-119(+)]; eri-1(mg366); rde-1(ne219); lin-15B(n744)*, XE1582: *wpSi11[eat-4p::rde-1::SL2::sid-1 + Cbr-unc-119(+)]; eri-1(mg366); rde-1(ne219); lin-15B(n744)*, N2 *[Ppfkb-1.1::GFP]*, N2 *[Ppfkb-1.1::pfkb-1.1::GFP]*, CB1370: *daf-2(e1370)*, CF1038: *daf-16(mu86)*, RB1828: *dgk-5(ok2366)*, VC1383: *dgk-5(gk691)*, RB1206: *rsks-1(ok1255)*, TG38: *aak-2(gt33)*, RB2325: *sens-1(ok3157)*, *sesn-1(tm2872)*, TJ356: *zIs356[daf-16p::daf-16a/b::GFP + rol-6(su1006)]*, CF1553: *muIs84 [sod-3p::GFP + rol-6(su1006)]*, N2 *[pitp-1p::GFP; myo-2p::mRFP]*, N2 *[pitp-1p::pitp-1::GFP; myo-2p::mRFP]*, N2 *[pitp-1p::pitp-1; myo-2p::mRFP]*. Strains were obtained from *Caenorhabditis* Genetics Center (CGC) and National BioResource Project (NBRP), unless noted otherwise. TU3401, TU3311, VP303, WM118, TJ356, RB2325 and *sesn-1(tm2872)* were provided by Dr. Chang-Shi Chen (National Cheng Kung University, Taiwan). XE1375, XE1474, XE1581, XE1582 were provided by Dr. Tsui-Ting Ching (National Yang Ming Chiao Tung University, Taiwan). For generation of N2 *[pitp-1p::GFP; myo-2p::mRFP]*, N2 *[pitp-1p::pitp-1::GFP; myo-2p::mRFP]* and N2 *[pitp-1p::pitp-1; myo-2p::mRFP]*, a plasmid DNA mix consisting of 80 ng/µL of *pitp-1* constructs and 20 ng/µL of co-injection marker, [*myo-2p::mRFP*], were microinjected into the gonad of young adult N2 hermaphrodite animals. For generation of N2 [*pitp-1p::ptr-23; myo-2p::tdtomato*], a plasmid DNA mix consisting of 80 ng/µL of [*pitp-1p::ptr-23*] and 20 ng/µL of co-injection marker, [*myo-2p::tdtomato*], were microinjected into the gonad of young adult N2 hermaphrodite animals. Individual F2 progenies were isolated to establish independent lines. Microinjection of N2 worms with co-injection marker, [*myo-2p::mRFP*] or [*myo-2p::tdtomato*], alone did not affect the mean lifespan of wild-type animals when grown on OP50 or HT115(DE3) bacteria (data not shown).

### RNA interference assay

*E. coli* HT115(DE3) (*Caenorhabditis* Genetics Center, Cat#HT115) transformed with either empty vector (L4440) or plasmid expressing double-stranded RNA for desired gene were cultured at 37°C overnight in LB supplemented with 100 µg/ml ampicillin and 100 µg/ml tetracycline. Bacteria were seeded on nematode growth medium (NGM) plates containing 100 µg/ml ampicillin and 1 mM IPTG. The RNAi clones picked from Julie Ahringer’s library (Source BioScience) were confirmed by sequencing using M13 forward primer (5’-TGTAAAACGACGGCCAGT-3’). RNAi clones made in this paper were constructed by inserting the cDNA of genes into the L4440 vector. For whole life RNAi, synchronized L1 were transferred to RNAi plates at 20°C. For adult only RNAi, worms were transferred from L4440 plates to RNAi plates at late L4 to young adult stage. Similarly, for RNAi from different adult age, worms were transferred from L4440 plates to RNAi plates at desired adult age.

### Lifespan assay

The worms used for lifespan assays were well fed and maintained at 20°C for at least three generations. All the lifespan assays were performed without using 5-fluoro-2’-deoxyuridine (FUdR). Synchronized worms were placed on NGM plates seeded with OP50 at 20°C. For RNAi conditions, synchronized worms were placed on NGM plates seeded with HT115(DE3) bacteria containing empty vector and were transferred to RNAi plates at desired age. For rapamycin treatment condition, rapamycin was dissolved in dimethyl sulfoxide (DMSO). The rapamycin solution was added into NGM plates with the final concentration of 100 µM rapamycin. The final concentration of DMSO for each plate was adjusted to 0.2%, including the control. All the plates used for treating rapamycin were used within 3 days. Synchronized worms were transferred from control plates to rapamycin plates at late L4 to young adult stage. When worms reached adulthood, worms were transferred to fresh plates with desired bacteria at a density of 25-35 worms per plate and were continually transferred to fresh plates every day until egg-laying ceased. After day 8 of adulthood, living worms were scored every 1-2 days and transferred to fresh plate every 3-7 days until all the worms were dead. Worms which did not move and did not respond to gently touch by platinum picker were scored as dead. Worms which exploded, crawled off plates, bagged or were accidentally killed were censored. Statistical analysis of lifespan data was performed by OASIS 2 [19].

### RNA extraction and quantitative real-time PCR

Synchronized worms were harvested at the desired stage and total RNA was extracted with REzol (Protech, PT-KP200CT). Worms were lysed by three times freezing and thawing by liquid nitrogen. RNA was purified by adding chloroform (Sigma, C2432) and precipitated by adding isopropanol (VWR, 0918). Extracted RNA was further purified by RQ1 RNase-Free DNase (Promega, # M6101). 1 µg of total RNA added with random primer (Promega, C118A) and M-MLV reverse transcriptase (Promega, M1701) was used for synthesis of cDNA according to the manufacturer’s instructions. Quantitative real-time PCR was set up by using *Power* SYBR™ Green PCR master mix (ABI, 4367659) and performed the reactions by ABI StepOnePlus Real-Time PCR System. The relative expression levels were calculated by ΔΔCt which was normalized by the internal control, *act-1*. The *p*-values were calculated by unpaired student t-test. qPCR primers used in this study are listed below. *act-1*, F: 5’-CGCCAACACTGTTCTTTCCG-3’; R: 5’-CTTGATCTTCATGGTTGATGGGG-3’. *pitp-1*, F: 5’-GGACAAGGTTCAAGATCGCC-3’; R: 5’-CTCACGGGAAAGAGCAACCA-3’. *Y54F10AR.1*, F: 5’-CATCCAGACTCCACTCC-3’; R: 5’-CGTATGCGCGTAGTTTTCGAC-3’. *Y71G12B.17*, F: 5’-GGTCTCCTATACGCAGTGTCG-3’; R: 5’-CGGACGCAGTGTGTTACTTG-3’. *sod-3*, F: 5’-GGGAGCACGCCTACTACTTG-3’; R: 5’-AGCATTGGCAAATCTCTCGC-3’. *dod-24*, F: 5’-TGTCCAACACAACCTGCATT-3’; R: 5’-TGTGTCCCGAGTAACAACCA-3’.

### Body bending assay

Synchronized worms at the desired stage were harvested from the NGM-OP50 agar plate to the M9 buffer. Each worm was allowed to adapt to the environment for 1 min, and the number of body bends was counted in 30 seconds under stereomicroscope. A body bend was considered as a change of direction of the cephalic region of a worm, denoted by the presence of the pharyngeal bulb towards the right side. Bending rates was calculated as body bend per second. The *p*-values were calculated by Two-way ANOVA.

### Paralysis assay

Paralysis assay were performed on fresh NGM plates. Worms at day 14 adulthood were placed on NGM plates and gently touched by platinum-made picker several times. If worms cannot escape from original location but still can move their head or pump their pharynx, these worms were considered as paralyzed worms. Worms that can escape form original location after touch or dead worms were not counted as paralyzed worms. The *p*-values were calculated by One-way ANOVA or unpaired student t-test.

### Body size measurement

Synchronized worms at the desired stage were harvested from the NGM-OP50 agar plate to 2% agarose pad. Photos were taken with a CCD camera (Nikon DS-Ri2) attached to a stereoscopic microscope (Nikon SMZ1500). Open Lab ver.2.2.5 software (Improvision) were used to measure the body size of each worm. The *p*-values were calculated by One-way ANOVA.

### Oxidative stress assay

The oxidative stress assay was conducted at 20°C. Young adult hermaphrodites were transferred to 160 µM or 240 µM juglone (5-hydroxyl-1,4-naphthoquinone, sigma, 481-39-0) containing NGM plates which were seeded with bacteria but without adding FUdR to induce oxidative stress. The number of dead worms was recorded every 2-6 hours until all worms were dead or censored from analysis because of worms exploded, crawled off plates, bagged or accidentally killed. The survival curves were performed by the percentage of death and the p-values were calculated by log-rank test. Statistical analysis of lifespan data was performed by OASIS 2 [19].

### Gene Expression Omnibus (GEO) analysis

Gene Expression Omnibus database (http://www.ncbi.nlm.nih.gov/geo) is an open functional genomics database of high-throughput resource. In this study, we downloaded the microarray data of GSE21784, GSE53890, GSE106672 and GSE77109 from GEO database. The microarray data of GSE21784 contained three biological repeats of synchronized populations of *C. elegans* at three points during aging. The microarray data of GSE53890 contained several samples of adult human brain samples from frontal cortical regions at different ages. The microarray data of GSE106672 contained four biological repeats of synchronized N2 (Bristol), *daf-2(e1370)*. The microarray data of GSE77109 contained 2 replicates, each was collected on day 4 of adulthood, fed by HT1115 bacteria. Gene expression was analyzed by GEO2R to compare two or more groups of samples. The *p*-values were calculated by unpaired student t-test.

### Western blot

Synchronized worms were harvested at the desired stage. 300-500 worms per sample were washed three times by M9 buffer and collected into 1.5mL tubes. After removing supernatants, WCE buffer (20 mM HEPES, pH 7.4, 0.2 M NaCl, 0.5 % Triton X-100, 5% glycerol, 1 mM EDTA, 10 mM β−glycerophosphate, 2 mM NA3VO4, 1mM NaF, 1 mM DTT) with 1x cocktail protease inhibitor (Roche) and 1x phosphatase inhibitor (Roche) was added to each sample. Samples were added with 0.5 mm ZrO beads and homogenized by the Bullet Blender. The concentration of extracted supernatants was detected by Bradford protein assay. The quantified proteins were mixed with 6X sample buffer dye (100 mM tris-HCl, pH 6.8, 4% SDS, 0.2% bromophenol blue, 200 mM 2-mercaptoethanol, 20% glycerol, 8M urea) and denatured at 95°C. 30 µg proteins were loaded in 10% SDS-PAGE for protein electrophoresis and transferred to NC-membrane by Bio-Rad system. The NC membrane was incubated in 5% BSA in 1x TBST as the blocking buffer. Immunoblotting was performed by incubating with anti-pS6K (Cell Signaling, Billerica, MA, USA, #9209, 1:500 dilution in 5% BSA /1xTBST), anti-p-AKT (Cell Signaling, #9271, 1:1000 dilution in 5% BSA/1xTBST), anti-puromycin (Merck Millipore, #MABE343, 1:5000 in 5% milk/1XTBST), anti-β-actin (GeneTex, GTX109639, 1:10 000 dilution in 5% milk/1xTBST), or anti-GAPDH (Epitomics, #S0011, 1:2000 in 5% milk/1XTBST). The membrane was washed three times with 1xTBST and incubated with the secondary antibody (Peroxidase-conjugated AffiniPure Goat Anti-Rabbit IgG (H+L), Jackson, 111-035-003, 1:10000 in 5 % BSA/1xTBST for phosphorylated proteins or 5 % milk/1xTBST for other proteins). After three times washing by 1xTBST, membrane was incubated with chemiluminescent HRP substrate (Millipore, WBKLS0500) and detected the chemiluminescent signals by ImageQuant LAS 4000 mini. The protein image was quantified by Image J to calculate the fold changes by normalizing each measurement to its control. The *p*-values were calculated by unpaired student t-test.

### Puromycin incorporation assay

To evaluate global protein synthesis in *C. elegans*, we employed a puromycin incorporation assay adapted with modifications from previously published protocols [20]. Synchronized worms were aged to day 5 of adulthood and collected using M9 buffer. Approximately 500 animals were washed twice with M9 and then resuspended in S-basal medium. For puromycin treatment, OP50 bacteria were grown overnight and subsequently concentrated 10-fold in S-basal. Worms were incubated in a 1 mL mixture composed of 750 µL S-basal, 200 µL of the concentrated OP50 suspension, and 50 µL of 10 mg/mL puromycin (Sigma, SI-P8833), yielding a final puromycin concentration of 0.5 mg/mL. Worms were then incubated in a mixture at 200 rpm for 4 hours at room temperature. Following treatment, worms were washed three times with ice-cold S-basal, chilled on ice, and snap-frozen in liquid nitrogen. Protein lysates were prepared using RIPA buffer, and puromycin-labeled proteins were detected by anti-puromycin via Western blotting as described previously. After blot stripping, β-actin was probed and used as a loading control.

### SOD-3 and DAF-16 reporter assay

The transgenic strains TJ356 and CF1553 were used in this study. Synchronized worms fed with EV or RNAi clones were harvested at the desired stage. Photos were taken with a CCD camera (Nikon DS-Ri2) attached to a stereoscopic microscope (Nikon SMZ1500) with the X-Cite® 120Q excitation light source (excitation at 470 nm and emission at 535 nm). The mean fluorescence intensity was measured by ImageJ software (NIH). The *p*-values were calculated by unpaired student t-test.

### RNAseq

Synchronized worms were harvested to day 3 of adulthood. The extracted RNA samples were DNase treated and assigned RNA Integrity Number (RIN) quality control. The RNA sample with RIN>7.0 can be used to perform next generation RNA sequencing (150 bp, paired-end, ∼20 million reads/sample, ∼6G total). Gene expression level is measured by transcript abundance. HISAT2 software was used to read alignment and StringTie software was used to assemble RNA-Seq alignments into potential transcripts in this experiment. Differential expression analysis was performed using DEGseq2 software. |FoldChange| > 1.5 and q-value < 0.05 are taken as the differentially expressed gene screening standard. The gene ontology (GO) enrichment analysis and the KEGG pathway analysis were performed by DAVID (https://david.ncifcrf.gov/). The RNAseq raw data can be accessed by the GEO accession number GSE309580.

## Results

### Reduction of *pitp-1* Promotes Healthy Longevity in *C. elegans*

To examine whether *pitp-1* regulates lifespan in *C. elegans*, we obtained two different *pitp-1* mutants, *pitp-1(pe1297)* and *pitp-1(tm1500)*, for lifespan measurement. Both *pitp-1* mutants exhibit significant lifespan extension and lowered *pitp-1* mRNA levels compared to wild-type N2 worms (Fig. 1A, 1B and Supplementary Table S1). Similarly, RNAi knockdown of *pitp-1* from day-1 adult (D1A) stage significantly extends lifespan compared to the control worms fed with empty vector (*EV*) (Fig. 1C, 1D and Supplementary Table S1). These results indicate that reduction of *pitp-1* expression promotes longevity in *C. elegans*.

**Figure 1.**
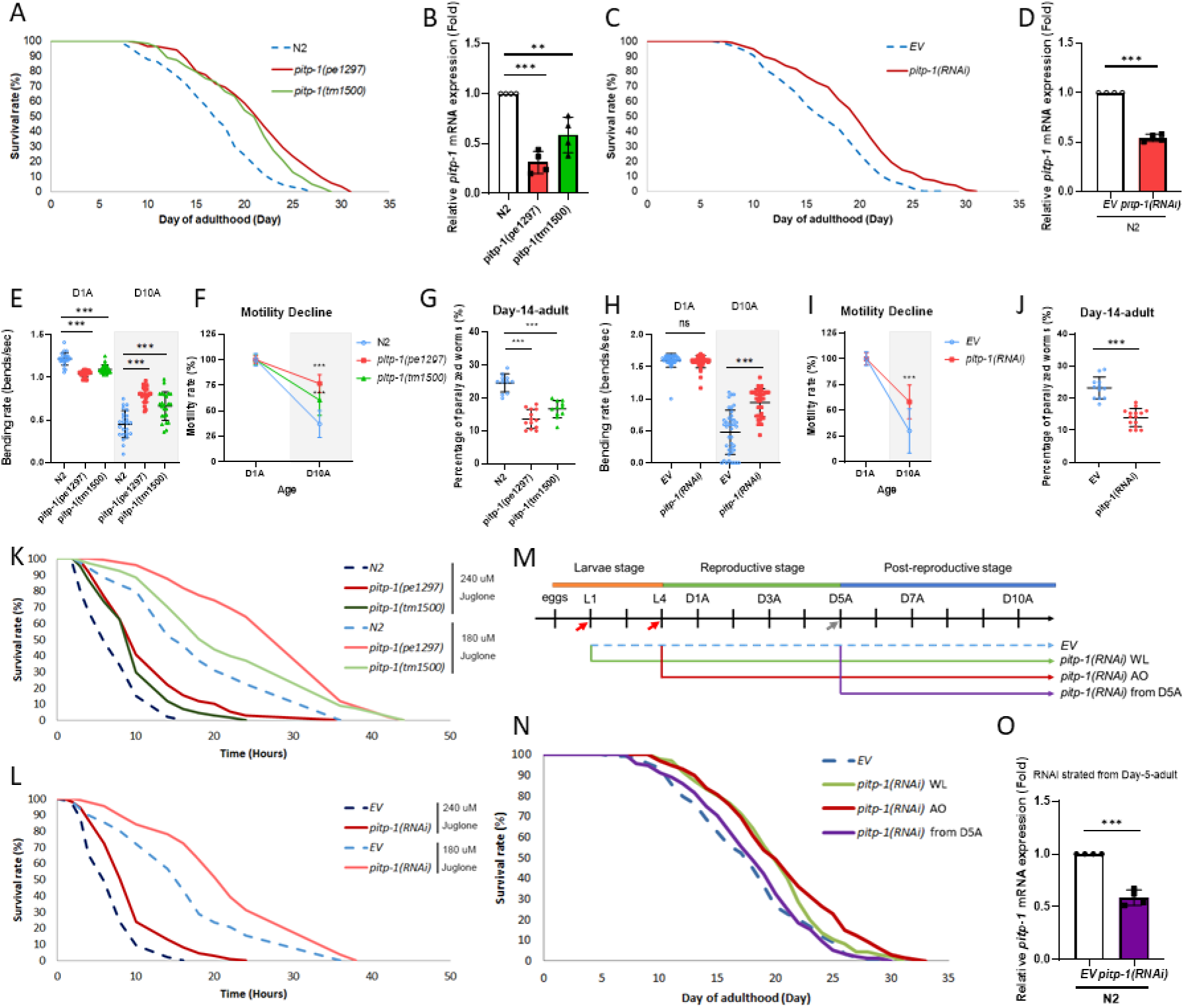
Reduction of *pitp-1* extends lifespan and promotes healthspan. (A) Two independent *pitp-1* mutants displayed significantly extended lifespan. (B) qPCR confirmed reduced *pitp-1* mRNA levels in *pitp-1* mutants. (C) Knockdown of *pitp-1* by RNAi from day-1 adult (D1A) extended lifespan. (D) qPCR confirmed reduced *pitp-1* mRNA expression upon *pitp-1(RNAi)*. (E, F) Both *pitp-1* mutants exhibited increased motility and ameliorated motility declines at D10A. (G) Both of the *pitp-1* mutants exhibited less paralyzed worms at D14A. (H, I) Knockdown of *pitp-1* increased motility and ameliorated motility declines at D10A. (J) Knockdown of *pitp-1* by RNAi ameliorated age-induced paralysis at D14A. (K, L) *pitp-1* mutants RNAi-treated worm showed enhanced resistance to juglone-induced oxidative stress. (M) Schematic diagram of RNAi treatment timeline. (N) *pitp-1* knockdown during adulthood extended lifespan. (O) qPCR confirmed that *pitp-1*(RNAi) from D5A reduces *pitp-1* mRNA levels. P-values were calculated by log-rank t-test in (A, C, K, L, N), and by One-way ANOVA in (B, F, G), and by Two-way ANOVA in (E, H), and by unpaired student’s t-test in (D, I, J, O). Survival curves are representative of three independent experiments, except oxidative stress assays, which were repeated twice with consistent results. Statistical significance was determined by log-rank test for lifespan assays, ANOVA for multiple comparisons, and unpaired Student’s t-test where applicable.

*pitp-1* is the single class II PITP gene with a conserved PITP domain in *C. elegans*. Given that the reduction of *pitp-1* extends lifespan, we wondered whether inhibition of the class I PITP genes, *Y54F10AR.1* or *Y71G12B.17*, would have a similar effect on lifespan. However, knockdown of *Y54F10AR.1* or *Y71G12B.17*, alone or in combination, did not prolong lifespan (Supplementary Fig. 1A and Supplementary Table S1). qPCR confirmed that *Y54F10AR.1* knockdown specifically reduced its own mRNA levels without affecting *Y71G12B.17* or *pitp-1* expression, and vice versa for *Y71G12B.17* knockdown (Supplementary Fig. 1B–D). These results suggest that only the reduced expression of class II PITP gene *pitp-1* plays a key role in promoting longevity in *C. elegans*.

We next examined whether reduction of *pitp-1* confers benefits on healthspan. Aging associates with motility decline and often culminates in paralysis. Thus, we measured locomotor capacity by quantifying body bending rates at day 1 and day 10 of adulthood (D1A and D10A). While *pitp-1* mutants exhibited slightly reduced bending rates compared to N2 at D1A, these mutants retained markedly higher bending rates than age-matched N2 at D10A (Fig. 1E, 1F), indicating the mutants do not display early-life hyperactivity but shows improved preservation of motility upon aging. RNAi knockdown of *pitp-1* produced similar benefits (Fig. 1H, 1I). Additionally, we assessed age-associated paralysis at D14A as another indicator of motor function. Paralysis at D14A was significantly reduced in both mutants and RNAi-treated worms (Fig. 1G, 1J), demonstrating a marked delay in paralysis onset. We next examined oxidative stress tolerance, another longevity-associated phenotype. We challenged both *pitp-1* mutants and RNAi-treated worms with juglone (5-hydroxy-1,4-naphthoquinone), a pro-oxidant compound that induces intracellular oxidative damage, and found both mutants and the RNAi-treated worms exhibited significantly increased survival (Fig. 1K, 1L and Supplementary Table S2). Long-lived organisms frequently display smaller body size. Consistent with this notion, both *pitp-1* mutants were significantly smaller than wild-type animals (Supplementary Fig. 1E). Together, these results demonstrate that reduction of *pitp-1* accompanies with multiple longevity-associated phenotypes, including improved motility, less paralysis, enhanced oxidative stress resistance, and reduced body size, supporting its role in promoting healthspan.

### Knockdown of *pitp-1* before post-reproductive age is essential for enhanced longevity

The timing of longevity intervention is critical, as different lifespan-regulating pathways show distinct temporal requirements. For example, in *C. elegans*, knockdown of *daf-2* during reproductive adulthood is important to extend lifespan [21]. Similarly, overexpression of *dFOXO* during reproductive adulthood promotes longevity in *Drosophila* [22]. In addition, the geroprotective benefits of TOR inhibition via long-term rapamycin treatment can be achieved with a short-term exposure to rapamycin during early adulthood in *Drosophila* and mice [23]. These data suggest the importance of investigating the precise time window for longevity assurance.

To delineate the temporal requirement of *pitp-1* reduction for extended lifespan, we initiated *pitp-1* RNAi knockdown from different stages: L1 stage, late L4 stage (adult-only, AO), and D5A (Fig. 1M). Knockdown of *pitp-1* from L1 or AO both significantly extended lifespan similarly, suggesting that *pitp-1* reduction from L1 larval development is dispensable and from AO is sufficient for promoting longevity (Fig. 1N and Supplementary Table S1). In contrast, *pitp-1* RNAi initiated at D5A with effective *pitp-1* mRNA reduction produced only a marginal increase for lifespan without significance (Fig. 1N, 1O), indicating a temporal restriction for the longevity effect. To further narrow the time window, we treated worms with *pitp-1* RNAi starting from D3A, D4A, D5A, or D7A (Supplementary Fig. 1F, 1G and Supplementary Table S1). *pitp-1* knockdown from D3A or D4A significantly extended lifespan, whereas *pitp-1* knockdown initiated at D5A or later failed to prolong lifespan (Fig. 1N; Supplementary Fig. 1G and Supplementary Table S1). Interestingly, the longevity effect of *pitp-1* knockdown was diminished when RNAi treated from D4A, suggesting that an optimal time window from L4 to D3A during early reproductive age. A previous microarray revealed that *pitp-1* expression declines with age in *C. elegans* [24], with significantly lower levels at D6A and D15A compared to L4 (Supplementary Fig. 2A). This age-associated *pitp-1* downregulation may explain why *pitp-1* knockdown from post-reproductive age no longer influences lifespan. Interestingly, analysis of human microarray data revealed similar age-dependent expression changes in *pitp-1* human orthologs [25]. Specifically, *PITPNM2* and *PITPNM3* expression was significantly reduced in the prefrontal cortex of extremely old individuals (>90 years) compared to younger adults (<40 years) (Supplementary Fig. 2B–2E), whereas *PITPNM1* showed a slight, non-significant decline (Supplementary Fig. 2F–2G). Together, these findings suggest that the longevity effect of *pitp-1* suppression is temporally restricted to a critical window prior to the post-reproductive stage, and that age-related reduction of *pitp-1*/PITPNMs expression may represent a conserved, protective feature of aging.

**Figure 2.**
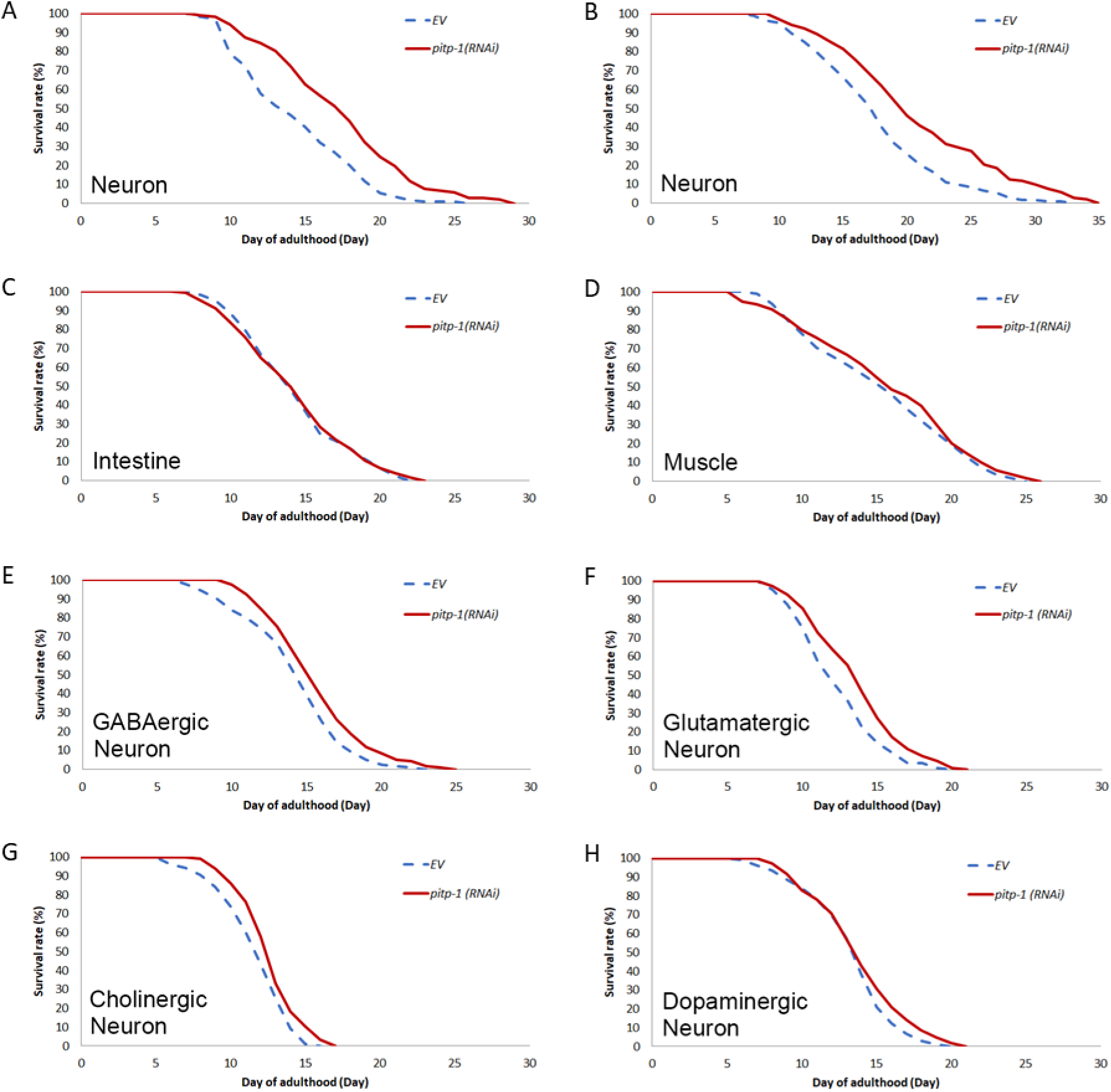
Reduction of *pitp-1* in pan-neuronal tissue and specific neuronal circuits extends lifespan. (A, B) Neuron-specific knockdown of *pitp-1* in *TU3401* and *TU3311* from adulthood increased lifespan. (C, D) Intestine-specific or Muscle-specific knockdown of *pitp-1* from adulthood did not alter lifespan. (E–G) Knockdown of *pitp-1* specifically in GABAergic neuron (*XE1375*), in glutamatergic neuron (*XE1582*), or in cholinergic neuron (*XE1581*) from adulthood extended lifespan. (H) Knockdown of *pitp-1* specifically in dopaminergic neuron (*XE1474*) showed no significant lifespan increase. Survival curves are representative of three independent experiments. Statistical significance was determined by log-rank test.

### The reduction of neuronal *pitp-1* is critical for lifespan extension

In addition to the temporal aspect, the spatial effect also plays an important role on longevity. For instance, increased neuronal or intestinal, but not muscular, DAF-16 activity is sufficient for lifespan extension in *C. elegans* [26]. In addition, neuronal TORC1 is essential for TOR-mediated aging regulation [27]. To evaluate the effect of tissue-specific *pitp-1* reduction on longevity, we performed RNAi knockdown in various tissue-specific RNAi strains. Neuronal knockdown of *pitp-1* significantly extended lifespan (Fig. 2A, 2B and Supplementary Table S3), whereas *pitp-1* knockdown in the intestine or muscle had no effect (Fig. 2C, 2D and Supplementary Table S3), indicating neuron is a crucial tissue for *pitp-1*-mediated lifespan regulation. To further investigate which neuron circuits may participate in the longevity upon *pitp-1* reduction, we performed *pitp-1* knockdown in either GABAergic, glutamatergic, cholinergic or dopaminergic neuronal circuit-specific strains individually for the lifespan measurement. Interestingly, except for dopaminergic neurons, knockdown of *pitp-1* in either GABAergic, glutamatergic, or cholinergic neurons is sufficient to extend lifespan (Fig. 2E-2H and Supplementary Table S3). These findings indicate that neuronal *pitp-1* suppression, particularly in certain specific neuron types, plays a central role in mediating its longevity effect.

### Overexpression of *pitp-1* decreases lifespan and impairs healthspan

To examine whether increased *pitp-1* has the opposite effects, we generated *pitp-1* overexpressing transgenic worms, one line with *pitp-1*::GFP fusion construct under *pitp-1* promoter, N2*[pitp-1p::pitp-1::GFP; myo-2p::mRFP]* (named N2 PITP-1 OE 1) with the control line, N2*[pitp-1p::GFP]* (named N2 control). To exclude possible GFP effects, we also generated *pitp-1* overexpressing without GFP fusion transgenic worms, N2*[pitp-1p::pitp-1; myo-2p::mRFP]* (named N2 PITP-1 OE 2). Confocal imaging confirmed the neuronal expression of the *pitp-1::GFP* fusion protein (Fig. 3A), consistent with previous findings[13]. Both strains showed about 3-4-fold increase in *pitp-1* mRNA (Fig. 3B). Opposite to *pitp-1* reduction, both N2 PITP-1 OE lines exhibited significantly shortened lifespan (Fig. 3C and Supplementary Table S1), reduced motility by at D10A (Fig. 3D), and increased paralysis at D14A (Fig. 3E), indicating deteriorated aging and health. Importantly, lifespan shortening in N2 PITP-1 OE worms was fully rescued by *pitp-1(RNAi)* (Fig. 3F), and overexpression of *pitp-1* in *pitp-1* mutants abolished their extended lifespan (Fig. 3G and Supplementary Table S4). These findings demonstrate that *pitp-1* acts as a negative regulator of healthy longevity in *C. elegans*.

**Figure 3.**
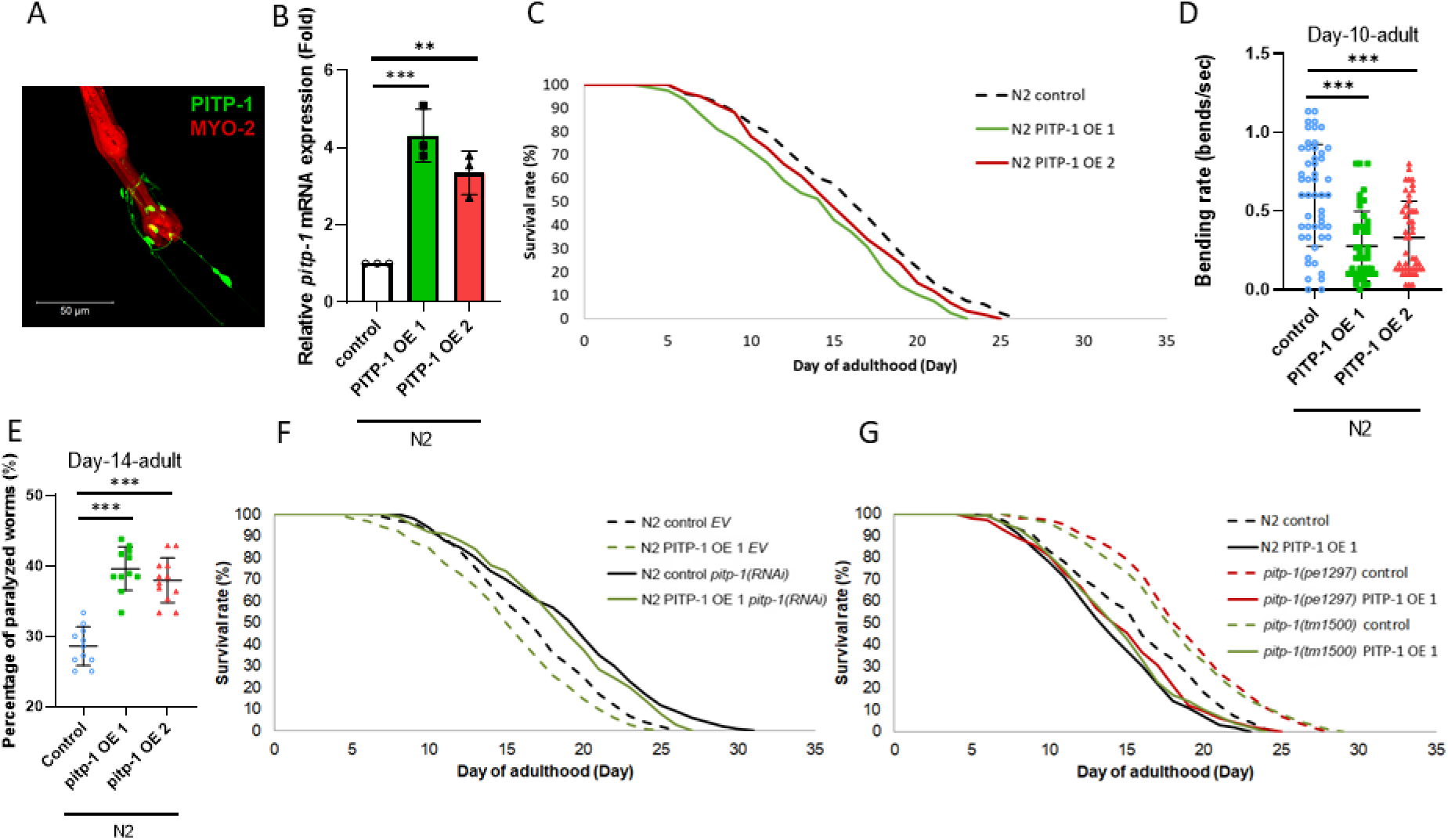
Overexpression of *pitp-1* decreases lifespan and impairs healthspan. (A) Confocal images showing *pitp-1* expression (GFP) and co-injection marker (mRFP) in PITP-1 OE 1. (B) qPCR confirmed elevated *pitp-1* mRNA levels in the *pitp-1* overexpression strains. (C) Overexpression of *pitp-1* reduced lifespan in both transgenic lines. (D, E) PITP-1-overexpressing worms displayed decreased body bending rate at D10A and increased paralysis at D14A compared to control line. (F) The reduced lifespan in N2 PITP-1 OE 1 can be rescued by *pitp-1*(RNAi) knockdown. (G) Overexpression of PITP-1 OE 1 reverted the extended lifespan in both *pitp-1* mutants. Survival curves are representative of three independent experiments. Statistical significance was determined by log-rank test for lifespan assays, ANOVA for multiple comparisons.

### *pitp-1* negatively regulates lifespan through modulating TOR signaling

Our previous study demonstrated that reduced *dgk-5* extends lifespan through downregulation of TOR signaling [6]. Since *pitp-1* and *dgk-5* function in the same pathway and that reduced expression of either gene leads to longevity, we hypothesized that *pitp-1* may also regulate lifespan via TOR signaling. Supporting this notion, knockdown of *pitp-1* did not further enhance the extended lifespan of *dgk-5* mutants (Supplementary Fig. 3A, 3B and Supplementary Table S4), suggesting *pitp-1* and *dgk-5* act through a common mechanism. Moreover, both *pitp-1* mutants and RNAi-treated worms exhibited significantly reduced p-S6K levels (Fig. 4A-4D), indicating diminished TOR signaling. In contrast, overexpression of *pitp-1* markedly increased p-S6K abundance (Fig. 4E and 4F), suggesting enhanced TOR activation. Furthermore, the elevated TOR signaling in *pitp-1*-overexpressing worms was suppressed by RNAi targeting *let-363*/TOR or its upstream activator *raga-1* (Supplementary Fig. 3C and 3D). Re-expression of *pitp-1* in the *pitp-1* mutant background restored p-S6K levels (Supplementary Fig. 3E, 3F), supporting a role for *pitp-1* as an upstream positive regulator of TOR signaling. Since TOR activation promotes protein translation, we further examined the effect of *pitp-1* on global translation. Both *pitp-1* mutants and RNAi-treated animals showed significantly reduced puromycin labeling (Fig. 4G–4J), indicating decreased protein translation. Conversely, overexpression of *pitp-1* markedly enhanced puromycin incorporation (Fig. 4K, 4L), representing elevated translational output. These results support that *pitp-1* positively regulates TOR activity and downstream protein synthesis. Furthermore, the extended lifespan of *pitp-1* mutants was not further prolonged by either genetic or pharmacological inhibition of TOR (Fig. 4M, 4N, Supplementary Fig. 3G, 3H and Supplementary Table S4). Similarly, *pitp-1* RNAi in the *rsks-1/S6K* mutant background failed to further extend the prolonged lifespan (Supplementary Fig. 3I and Supplementary Table S4). Moreover, suppression of TOR signaling by either *let-363*(RNAi) or rapamycin treatment rescued the lifespan shortening and reduced motility caused by *pitp-1* overexpression (Fig. 4O, 4P, Supplementary Fig. 3J and Supplementary Table S4). Collectively, these results demonstrate that *pitp-1* negatively regulates lifespan through modulation of TOR signaling.

**Figure 4.**
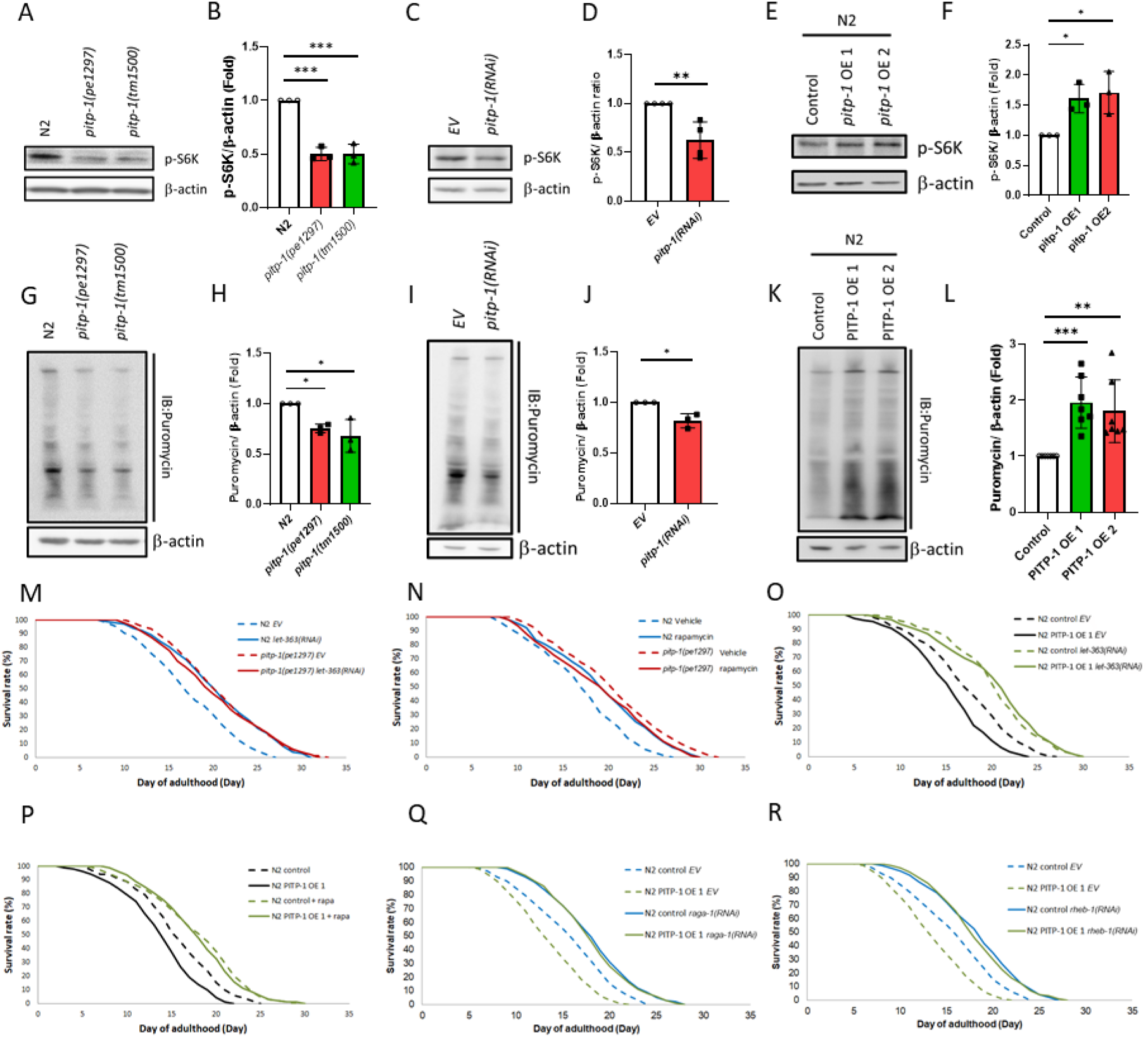
*pitp-1* negatively regulates lifespan through modulating TOR signaling. (A–F) *pitp-1* mutants or RNAi knockdown reduced phosphorylated S6K levels, whereas PITP-1 overexpression increased S6K phosphorylation. (G–L) puromycin incorporation assays showed reduced protein synthesis in *pitp-1* mutants or RNAi animals and elevated levels in PITP-1-overexpressing strains. (M, N) Genetic or pharmacological inhibition of TOR by *let-363*(RNAi) or rapamycin did not further extend the longevity of *pitp-1* mutants. (O, P) TOR inhibition rescued the shortened lifespan caused by PITP-1 overexpression. (Q, R) Knockdown of upstream TOR regulators *raga-1* or *rheb-1* rescued the reduced lifespan of PITP-1-overexpressing worms. Survival curves are representative of three independent experiments. Statistical significance was assessed by log-rank test for lifespan assays, ANOVA for multiple comparisons, and unpaired Student’s t-test where applicable.

### Several TOR regulators are involved in *pitp-1*-mediated lifespan regulation

TOR activity is modulated by amino acids, growth factors, and energy stress via distinct regulators (Supplementary Fig. 3K). To identify upstream regulators linking *pitp-1* to TOR activity, we performed lifespan epistasis tests. RNAi knockdown of *raga-1* or *rheb-1* rescued the shortened lifespan in *pitp-1*-overexpressing animals (Fig. 4Q, 4R and Supplementary Table S4), suggesting that both Rag and Rheb GTPases are involved in *pitp-1*-mediated lifespan regulation. Conversely, knockdown of *pitp-1* still extended lifespan in the AMPK-deficient strain *aak-2(gt33)* (Supplementary Fig. 3L and Supplementary Table S4), suggesting that *pitp-1* regulates lifespan independent of AMPK. Sestrin, a negative regulator of Rag GTPases, is known to inhibit the amino acid sensing arm of TORC1 and promotes longevity in *C. elegans* [28]. Given the role of Rag GTPases in *pitp-1*-mediated lifespan regulation, we next investigated whether *sestrin* is also required for this effect. Notably, the lifespan extension induced by *pitp-1* knockdown was abolished in *sesn-1* mutant animals (Supplementary Fig. 3M, 3N and Supplementary Table S4), indicating that *sestrin* is required for *pitp-1*-mediated lifespan extension. Together, these findings reveal that the sestrin–Rag GTPase axis and Rheb GTPase, upstream regulators of TOR, are involved in *pitp-1*-mediated lifespan regulation.

### *pitp-1* is involved in insulin/IGF-1 signaling-mediated lifespan regulation

Because AKT lies upstream of Rheb-TOR, we first detected the p-AKT levels in long-lived *pitp-1* mutants to check IIS involvement in *pitp-1*-mediated lifespan regulation. Both *pitp-1* mutants exhibit significantly reduced p-AKT levels compared to N2 worms (Fig. 5A and 5B), suggesting that IIS may be involved in *pitp-1*-mediated lifespan extension. However, knockdown of *pitp-1* did not promote DAF-16::GFP nuclear translocation nor increase expression of DAF-16 target gene *sod-3* (Supplementary Fig. 4A-4C). These results indicate that *pitp-1* knockdown does not promote DAF-16 transcription activity. To further clarify if DAF-16 is required for *pitp-1* knockdown-mediated lifespan extension, we performed lifespan assays in the *daf-16(mu86)* null mutant by *pitp-1* knockdown. Knockdown of *pitp-1* still extended lifespan in *daf-16(mu86)* mutant (Fig. 5C and Supplementary Table S4), indicating that DAF-16 is not required for the longevity effect by *pitp-1* suppression.

**Figure 5.**
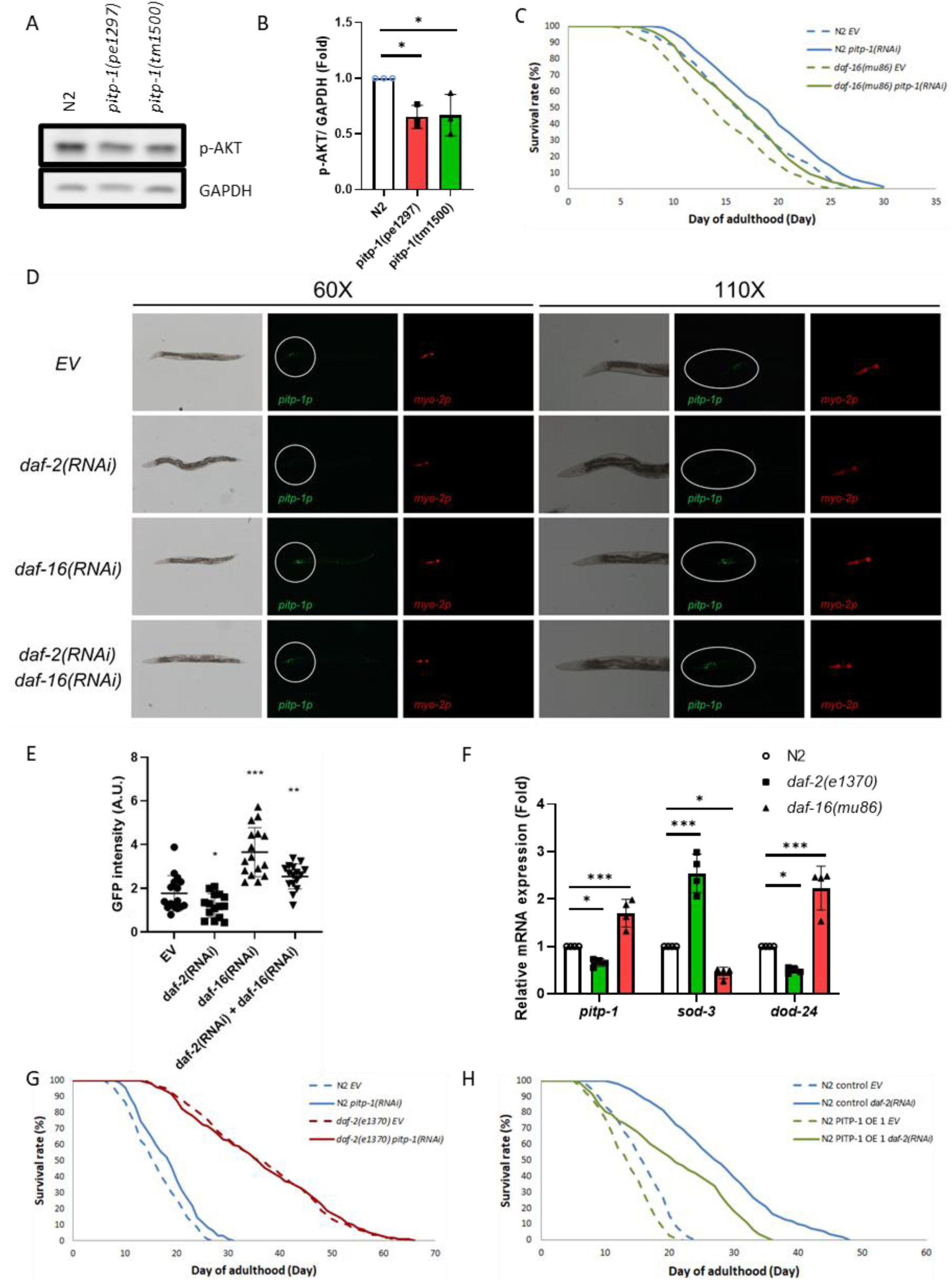
Integration of *pitp-1* in insulin/IGF-1 signaling–mediated lifespan regulation. (A, B) *pitp-1* mutants exhibited reduced p-AKT levels compared to N2. (C) Knockdown of *pitp-1* extended lifespan in *daf-16(mu86)*. (D–F) Transcriptional reporter and qPCR analyses showed that *pitp-1* expression is negatively regulated by IIS: *daf-2* knockdown reduced expression, while *daf-16* knockdown increased it. Known DAF-16 targets (*sod-3*, *dod-24*) were used as positive controls. (G) Knockdown of *pitp-1* did not further extend lifespan in *daf-2(e1370)*. (H) PITP-1 overexpression partially blocks the longevity effect by *daf-2(RNAi)* knockdown. Survival curves are representative of three independent experiments. Statistical significance was determined by log-rank test for lifespan assays, ANOVA for multiple comparisons.

Interestingly, two DAF-16 binding sites were identified in the *pitp-1* promoter [29], raising the possibility that *pitp-1* may act downstream to DAF-16 and be regulated by DAF-16. To test this, we analyzed *pitp-1* promoter-driven GFP expression following RNAi of *daf-2* or *daf-16*. Knockdown of *daf-2* significantly reduced *pitp-1*::GFP expression, while *daf-16* RNAi elevated its expression (Fig. 5D and 5E). Moreover, co-treatment with *daf-2* and *daf-16* RNAi restored the decreased *pitp-1* GFP intensity caused by *daf-2* knockdown (Fig. 5D and 5E). Similarly, qPCR confirmed that *pitp-1* transcript levels were downregulated in *daf-2(e1370)* but upregulated in *daf-16(mu86)* mutants (Fig. 5F). Consistently, a previous microarray study also reported reduced *pitp-1* expression in *daf-2(e1370)* (Supplementary Fig. 4D) [30]. These data indicate that *pitp-1* is transcriptionally repressed by DAF-16. Accordingly, *pitp-1* knockdown failed to further extend lifespan in *daf-2(e1370)* mutant (Fig. 5G and Supplementary Table S4), likely due to their already reduced *pitp-1* expression levels. Conversely, overexpression of *pitp-1* partially suppressed the lifespan extension induced by *daf-2(RNAi)* or *age-1(RNAi)* (Fig. 5H, Supplementary Fig. 4E and Supplementary Table S5), further supporting the notion that *pitp-1* acts as a downstream effector negatively regulated by IIS. Together, these results suggest that *pitp-1* functions downstream of DAF-16 and contributes to IIS-mediated lifespan regulation.

### Transcriptome-wide analyses uncover signaling shifts and longevity mechanisms upon *pitp-1* suppression

Since the longevity effect of *pitp-1* suppression is temporally restricted, and likely involves complex transcriptional reprogramming and pathway cross-talk, we performed RNA sequencing on D3A worms with reduced *pitp-1* expression to gain a comprehensive understanding of transcriptomic changes. Transcriptomic profiles were analyzed using Qiagen Ingenuity Pathway Analysis (IPA) and Over-Representation Analysis (ORA) (Fig. 6A). IPA canonical pathway analysis revealed downregulation of IIS, PI3K/AKT and TOR in *pitp-1* mutant and *pitp-1(RNAi)*-treated worms (Fig. 6B), consistent with our mechanism study findings. In addition, PTEN signaling, the negative regulator of IIS, was activated upon *pitp-1* suppression, supporting *pitp-1* reduction leads to attenuation of IIS. In contrast, AMPK signaling was not activated, which is consistent with our previous results showing that *pitp-1* reduction does not promote longevity through AMPK activation (Supplementary Fig. 3L). Similarly, IPA upstream regulator analysis suggests that the transcriptomic changes we observed may result from decreased upstream activity of mTOR or insulin signaling (Fig. 6C). Together, these results reinforce the critical roles of IIS and TOR signaling in mediating *pitp-1*-dependent longevity.

**Figure 6.**
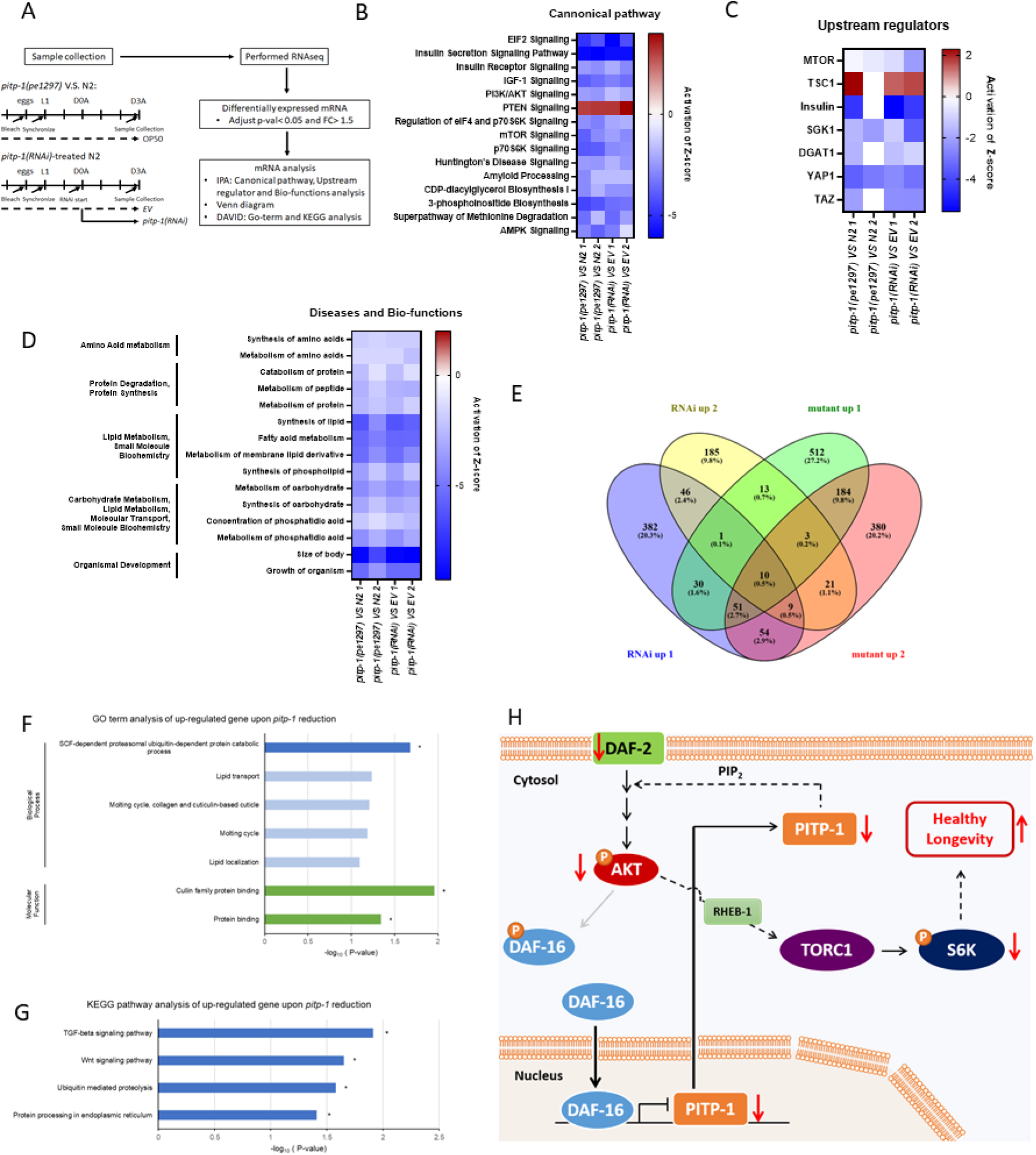
Transcriptomic and pathway analysis upon *pitp-1* reduction. RNA-seq and pathway enrichment analyses revealed downregulation of insulin/TOR signaling and upregulation of proteolysis-related genes in *pitp-1* mutants and RNAi-treated worms. (A) Schematic diagram of RNA-seq samples collection and data analysis. (B) The results of QIAGEN IPA canonical pathway analysis upon *pitp-1* reduction. (C) The results of QIAGEN IPA upstream regulator analysis upon *pitp-1* reduction. (D) The results of QIAGEN IPA disease and biofunctions analysis upon *pitp-1* reduction. (E) ORA analysis identified potential up-regulated downstream target genes which might involve in *pitp-1* reduction-mediated longevity. (F) GO terms enrichment analysis of the up-regulated genes upon *pitp-1* reduction (FC> 1.5; adjusted p<0.05*). The vertical coordinates were the enriched GO terms, and the horizontal coordinates were the numbers of the up-regulated genes in these GO terms. The blue columns represent the biological process GO terms. The green columns represent the molecular function GO terms. GO terms enrichment analysis was conducted by DAVID. (G) KEGG enrichment analysis of the changed genes upon *pitp-1* reduction (FC> 1.5; adjusted p<0.05*). KEGG enrichment analysis was conducted by DAVID. (H) The working model for *pitp-1* reduction-mediated lifespan regulation in *C. elegans*.

In addition to IIS and TOR, eIF2 signaling was also significantly downregulated in *pitp-1*-reduced worms (Fig. 6B). As a key regulator of translation initiation, inhibition of eIF2B enhances proteostasis and extends lifespan [20, 31]. Consistently, *pitp-1* suppression reduced protein synthesis (Fig. 4G-4J), accompanied by extended lifespan. Together, these observations suggest that *pitp-1* reduction promotes longevity, perhaps by improving protein homeostasis.

We also found that *pitp-1* suppression downregulated CDP-DAG biosynthesis pathway and 3-phosphoinositide biosynthesis (Fig. 6B), suggesting PPI cycle activity may be downregulated. Consistently, IPA Diseases and Bio-functions analysis (Fig. 6D) revealed decreased lipid metabolism, amino acid metabolism, and protein synthesis, likely reflecting TOR suppression. This metabolic reduction may also explain the downregulation of biofunctions such as “size of body” and “growth of organism” (Fig. 6D). These results align with our earlier observation that *pitp-1* mutants exhibit smaller body size (Supplementary Fig. 1E).

In addition to IPA, we employed Over-Representation Analysis (ORA) to identify downstream gene targets upon *pitp-1* reduction. Using a cutoff of >1.5-fold change and adjusted *p*-value < 0.05, we identified 10 genes significantly upregulated upon *pitp-1* suppression (Fig. 6E). GO and KEGG pathway analyses revealed that the “SCF-dependent, ubiquitin-mediated proteasomal protein catabolic process”, “ubiquitin-mediated proteolysis” and “Protein processing in endoplasmic reticulum” was significantly overrepresented (Fig. 6F, 6G). This result aligns with our IPA analysis showing reduced global protein synthesis (Fig. 6D) and our puromycin incorporation assay (Fig. 4G-4J). Given that enhancing proteolytic systems, including the ubiquitin–proteasome pathway, promotes longevity [32], our findings suggest that *pitp-1* reduction may promote healthy longevity by maintaining proteostasis.

In summary, these transcriptomic analyses reinforce that *pitp-1* reduction promotes longevity through coordinated suppression IIS, TOR signaling, reducing anabolic activity and enhancing proteostasis. Our data support a model in which DAF-16 represses *pitp-1* transcription under reduced IIS, partially contributing to IIS-mediated longevity. Reduced *pitp-1* attenuates TOR activity via the AKT–RHEB axis, thereby promoting longevity. This is the first study to identify *pitp-1* as a novel lifespan regulator and highlights its involvement in IIS–TOR crosstalk, offering new insights into aging regulation and potential anti-aging interventions.

## Discussion

PITP is a critical regulator in PPI turnover, but its role in aging remains unclear. In this study, we identify a novel function for *pitp-1*, a Class II PITP, in regulating lifespan and healthspan in *C. elegans*. We show that *pitp-1* is transcriptionally repressed by DAF-16 and acts as a pro-aging factor by promoting TOR signaling. Notably, our spatial and temporal analyses reveal that both neuronal specificity and early adulthood timing are essential for *pitp-1*-mediated lifespan regulation. These findings position *pitp-1* as a critical regulator that connects the IIS and TOR pathways in aging control, with its pro-aging function constrained by specific neuronal and temporal contexts.

Our temporal analysis highlighted a critical window during early adulthood, particularly the early reproductive stages, as essential for *pitp-1*-mediated lifespan regulation, with adult-onset knockdown sufficient to promote healthy longevity without interfering with development. This time window is consistent with the concept that interventions in nutrient-sensing pathways are most effective during this stage [21, 22, 23]. These parallels highlight that *pitp-1* reduction aligns with these conserved longevity-regulating pathways during a critical early-adult temporal window. Moreover, our analysis of public gene expression datasets revealed a natural, age-associated decline in *pitp-1/PITPNMs* expression in worms and humans (Supplementary Fig. 2) [24, 25], suggesting that this downregulation may represent a conserved protective mechanism against aging.

Spatially, neuronal knockdown of *pitp-1* is sufficient to promote longevity, emphasizing the central role of the nervous system in systemic aging regulation where IIS and TOR signaling exert their lifespan-modulating effects [26, 27, 33]. Consistently, neuronal inhibition of RAGA-1 from hatching or D1A extends lifespan, supporting the temporal flexibility of neuronal TOR suppression in promoting longevity [34]. In addition, other PPI cycle genes, *inaE/dagl/dagl-1* and *egl-8/PLCβ*, expressed in neurons also regulate lifespan through the TOR pathway [6, 35]. These findings reinforce neuronal modulation of PI signaling impacts systemic aging via TOR, in line with the role we propose for *pitp-1*.

Moreover, suppression of *pitp-1* in either glutamatergic, cholinergic, or GABAergic neurons each extended lifespan, suggesting that *pitp-1* exerts its pro-aging function through multiple excitatory and inhibitory neuronal circuits. Inhibition of age-related increases in neural excitation, particularly in glutamatergic and cholinergic neurons, extends lifespan [36]. Chronic hyperexcitability of glutamatergic neurons accelerates aging by PLCβ–IP3R pathway overactivation, and suppressing this pathway restores normal lifespan [36, 37]. Additionally, mTOR hyperactivation enhances synaptic responses in glutamatergic and GABAergic neurons, while rapamycin treatment normalizes glutamatergic overexcitation and restores neurotransmitter balance [38]. These data suggest that PLCβ–IP3R pathway or mTOR suppression in excitatory and inhibitory neurons promotes neural homeostasis and healthy aging, aligning with our findings. Taken together, *pitp-1* emerges as a neuronal regulator of longevity acting through TOR, potentially by modulating neuronal excitability and neurotransmission. Future studies should clarify the role of distinct circuits and how *pitp-1* coordinates PPI turnover to systemic metabolic responses upon aging.

eIF2 is a central regulator of translation initiation, whose activity is inhibited by phosphorylation of eIF2α, leading to global translational repression and proteostasis maintenance [20, 31]. Our IPA analysis revealed that *pitp-1* downregulation reduces eIF2 signaling, consistent with reduced global translation. Moreover, prior studies have reported bidirectional crosstalk between eIF2 and mTORC1: mTORC1 inhibition can activate GCN2 to phosphorylate eIF2α, whereas eIF2α phosphorylation and ATF4 translation can inhibit mTORC1 by REDD1 and Sestrin2 induction [39, 40]. In line with this, we found that *pitp-1* downregulation not only represses TOR signaling but also requires *sestrin* for lifespan extension (Supplementary Fig. 3M, 3N), suggesting the involvement of the eIF2α–ATF4–Sestrin axis. This pathway acts independently of AMPK [40], consistent with our data (Supplementary Fig. 3L), and may inhibit TOR by restraining Rag GTPase–mediated activation [28]. Together, our findings suggest that the observed downregulation of eIF2 signaling upon *pitp-1* reduction may potentially contribute to TOR inhibition and healthy longevity, while further studies are needed to establish a direct causal role in the future.

In this study, we also found that *pitp-1* suppression in *C. elegans* not only promotes longevity but also results in reduced body size, a phenotype often linked to altered nutrient signaling. In addition to reduced TOR or IIS activity, the two nutrient-sensing pathways known to influence body size, our transcriptomic analysis further revealed a downregulation of YAP and TAZ in *pitp-1*-suppressed worms, as predicted by upstream regulator analysis using IPA (Fig. 6C). This finding is consistent with recent studies in mammalian systems showing that inhibition of PITPα/β activates the Hippo pathway, leading to suppression of YAP-mediated transcription, reduced cell proliferation, and enhanced cancer cell death [41]. Our observation of reduced YAP/TAZ activity and smaller body size upon *pitp-1* suppression may reflect a conserved mechanism, where diminished PI4P-mediated suppression of the Hippo pathway contributes to reduced growth. Importantly, these findings raise the possibility that modulation of the Hippo pathway or direct regulation of its downstream transcription factors YAP/TAZ may represent a novel and promising strategy to promote healthy aging. In particular, investigating how *pitp-1* interfaces with the Hippo pathway may uncover a previously unrecognized lipid-signaling mechanism with relevance to both growth regulation and age-associated functional decline.

In addition to downregulation of canonical nutrient-sensing pathways, eIF2 signaling and Hippo pathway, our IPA analysis revealed a suppression of Huntington’s disease (HD) signaling upon *pitp-1* reduction (Fig. 6B). Given that mutated Huntingtin (Htt) enhances mTORC1 activity through Rheb interaction and aberrant PI3K/AKT/mTOR signaling contributes to HD pathogenesis [42], these findings are consistent with our model that *pitp-1* modulates lifespan through Rheb-TOR signaling and shows downregulation of HD-associated signaling. Notably, HD is characterized by dysfunction of both GABAergic and glutamatergic neurons, which are central to motor impairment and excitotoxicity [43]. Strikingly, suppression of *pitp-1* in either GABAergic or glutamatergic neurons was sufficient to extend lifespan in *C. elegans* (Fig. 2), suggesting that *pitp-1* may act through conserved neuronal circuits also implicated in HD pathogenesis. This raises the intriguing possibility that *pitp-1* suppression could not only promote healthy longevity but also provide therapeutic potential for neurodegenerative diseases such as HD. Future investigations exploring whether *pitp-1* modulation can mitigate HD-related phenotypes in mammalian models may uncover promising therapeutic avenues for neurodegenerative disease intervention.

Consistently, our parallel study in *Drosophila* revealed that downregulation of *rdgB*, the orthologue of *pitp-1*, also promotes healthy longevity and reduces TOR activity (data not shown). These findings strongly suggest that the pro-aging role of PITPs and their regulation of TOR signaling are evolutionarily conserved. Thus, the mechanisms uncovered here may extend beyond nematodes and flies, raising the exciting possibility that PITP modulation could exert similar effects on aging and healthspan in mammals, including humans.

## Conclusion

Our findings uncover *pitp-1* as a previously unappreciated regulator of aging, acting at the intersection of IIS and TOR signaling in a neuron- and age-specific manner. This study establishes *pitp-1* as a critical node coordinating nutrient-sensing pathways to regulate healthy longevity and highlights its potential as a target for aging-associated interventions. While our results reveal its role in IIS-TOR cross talk, it also suggest that *pitp-1* may influence additional pathways implicated in growth control, proteostasis, and neurodegeneration, including Hippo, eIF2, and Huntington’s disease–related signaling. These findings warrant further investigation, particular in other species such as *Drosophila* or mammals, future studies to assess the conservation of this regulatory axis and its relevance for both healthy aging and anti-neurodegenerative diseases.

## Data availability

All the RNAseq raw data can be accessed by the GEO accession number GSE309580.

## Abbreviations

pitp-1: phosphatidylinositol transfer protein-1
PITPs: phosphatidylinositol transfer proteins
PI: phosphatidylinositol
PA: phosphatidic acid
ER: endoplasmic reticulum
PM: plasma membrane
PPI: phosphoinositide
DAG: diacylglycerol
IIS: insulin/IGF-1 signaling
TOR: target of rapamycin
p-S6K: phosphorylated S6K
p-AKT: phosphorylated AKT
TORC1: TOR complex 1
TORC2: TOR complex 2
PLCβ: phospholipase C β
EV: empty vector
GEO: Gene Expression Omnibus
RNAseq: RNA sequencing
RIN: RNA Integrity Number
IPA: ingenuity pathway analysis
ORA: over-representation analysis
GO: gene ontology
CGC: Caenorhabditis Genetics Center
NBRP: National BioResource Pproject
NGM: nematode growth medium
FUdR: 5-fluoro-2’-deoxyuridine
DMSO: dimethyl sulfoxide
Juglone: 5-hydroxyl-1,4-naphthoquinone:
DTT: Dithiothreitol
NSTC: National Science and Technology Council

---

**Supplementary Figure 1.**
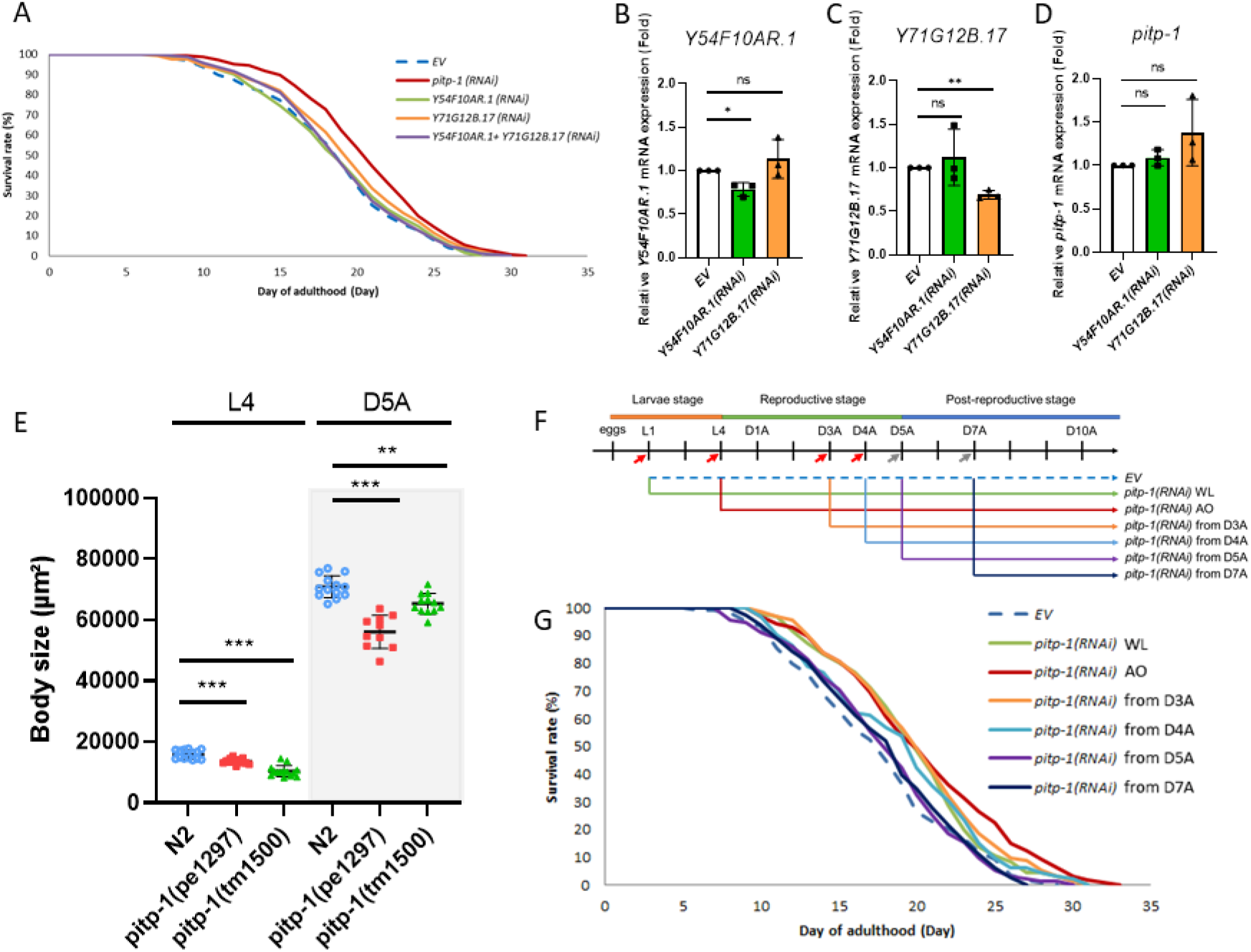
The reduction of class II PITP, *pitp-1*, reveals longevity-related phenotypes. (A) Knockdown of *pitp-1*, but not other class I PITP homologs, extended lifespan in N2. (B-D) qPCR confirmed RNAi against class I PITP homologs specifically reduced their own transcript levels without affecting *pitp-1*. (E) *pitp-1* mutants exhibited reduced body size. (F) Schematic diagram of RNAi treatment timelines. (G) *pitp-1* knockdown during the reproductive stage promotes longevity. Survival curves are representative of three independent experiments. Statistical significance was determined by log-rank test for lifespan assays, ANOVA for multiple comparisons.

**Supplementary Figure 2.**
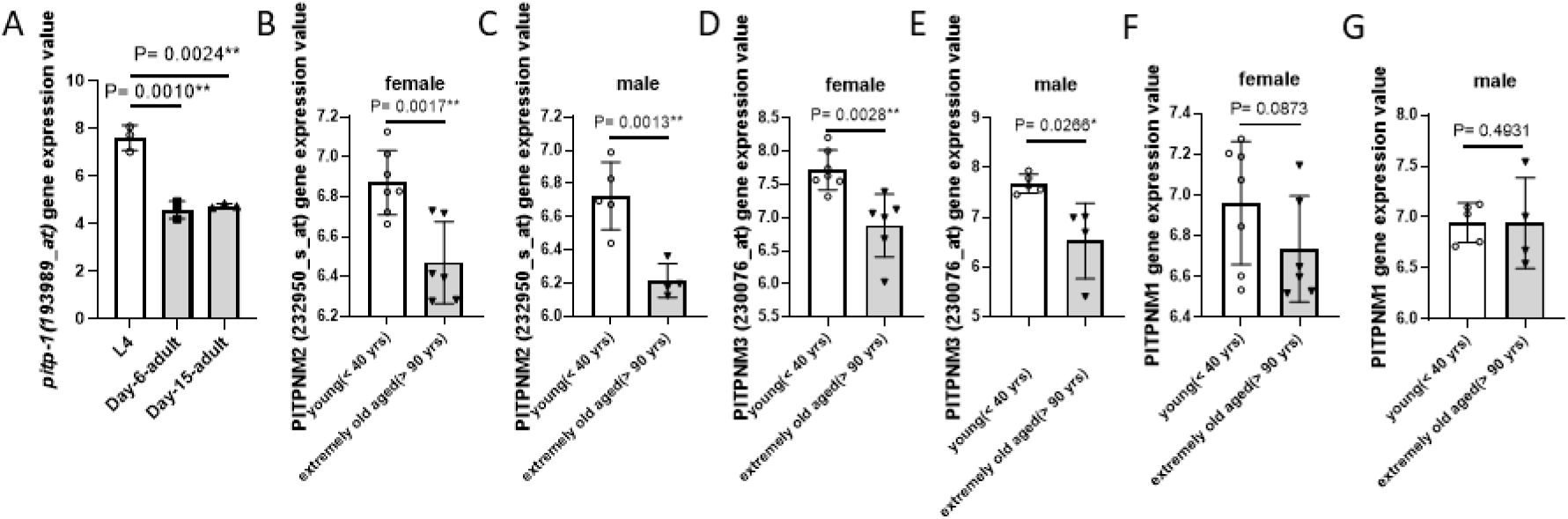
GEO shows reduced class II PITP expression in old age. (A) Whole-genome microarray data from *C. elegans* [24] revealed significant reductions in both *pitp-1* splice variants at day-6 and day-15 adults compared to L4 larvae (One-way ANOVA). (B-G) Microarray analysis of human frontal cortex [25] showed that expression of PITPNM2 (232950_at) and PITPNM3 (230076_at) was significantly lower in individuals >90 years (extremely old) compared to those <40 years (young) in both sexes, while PITPNM1 showed a slight, non-significant decrease (unpaired Student’s *t*-test).

**Supplementary Figure 3.**
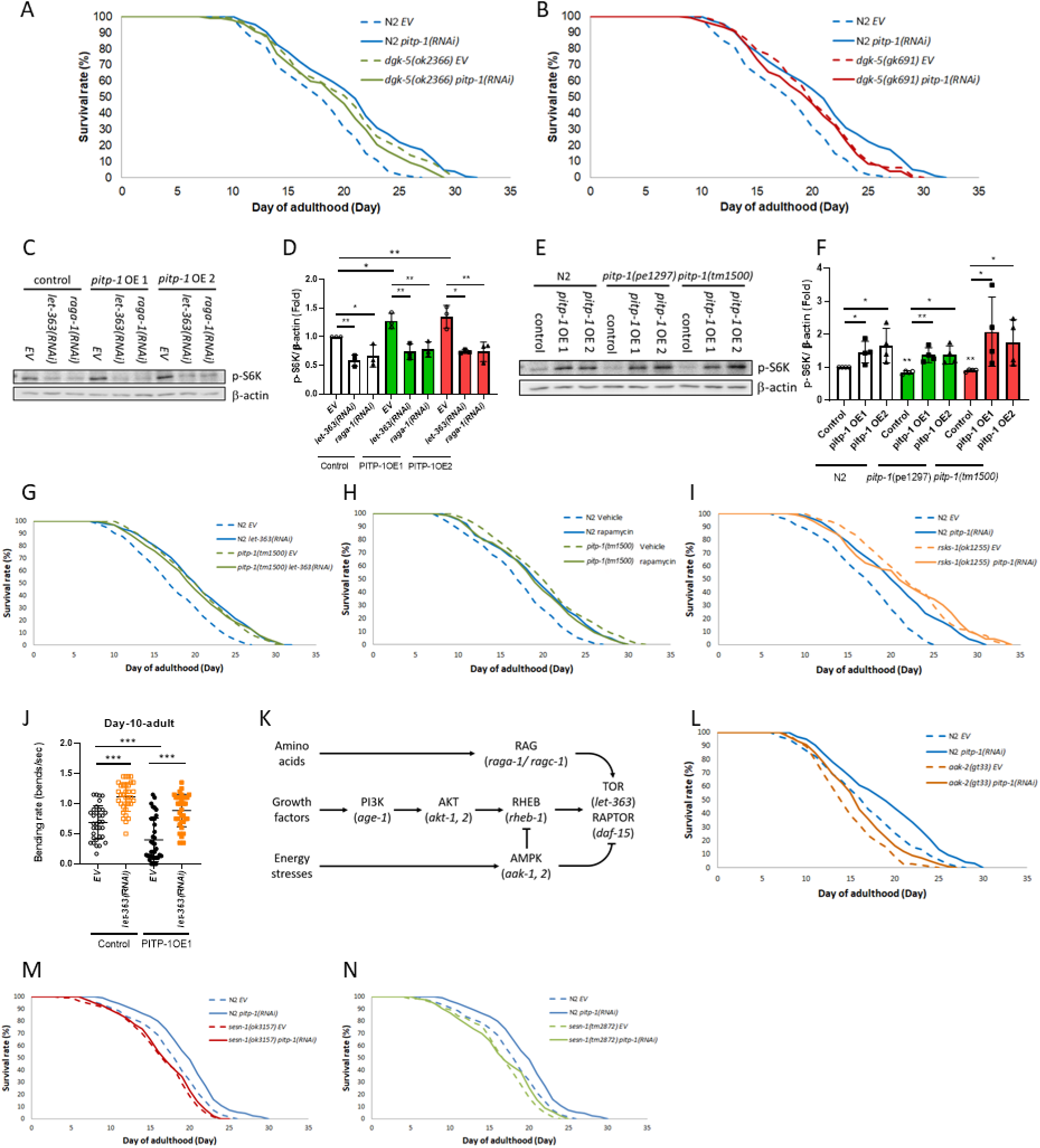
*pitp-1* negatively regulates lifespan by modulating TOR signaling. (A–B) RNAi knockdown of *pitp-1* did not further prolong the extended lifespan in two *dgk-5* mutants. (C, D) The elevated p-S6K levels in PITP-1 overexpression strains were reverted by genetic inhibition of TOR signaling (*let-363*, *raga-1*). (E, F) The reduced p-S6K levels in two *pitp-1* mutants were blocked by PITP-1 overexpression. (G, H) Genetic or pharmacological inhibition of TOR did not further enhance the extended lifespan in *pitp-1(tm1500)* mutant. (I) Knockdown of *pitp-1* did not further prolong the enhanced lifespan in *rsks-1* mutants. (J) Genetic knockdown of TOR by *let-363(RNAi)* rescued the motility defect caused by PITP-1 overexpression. (K) Schematic diagram of TOR upstream regulators RAG, RHEB, AMPK. (L) Knockdown of *pitp-1* extended lifespan in *aak-2(gt33)*. (M, N) *sesn-1* mutation blocked the longevity effect of *pitp-1(RNAi)* knockdown. Survival curves are representative of three independent experiments. Statistical significance was determined by log-rank test for lifespan assays, ANOVA for multiple comparisons.

**Supplementary Figure 4.**
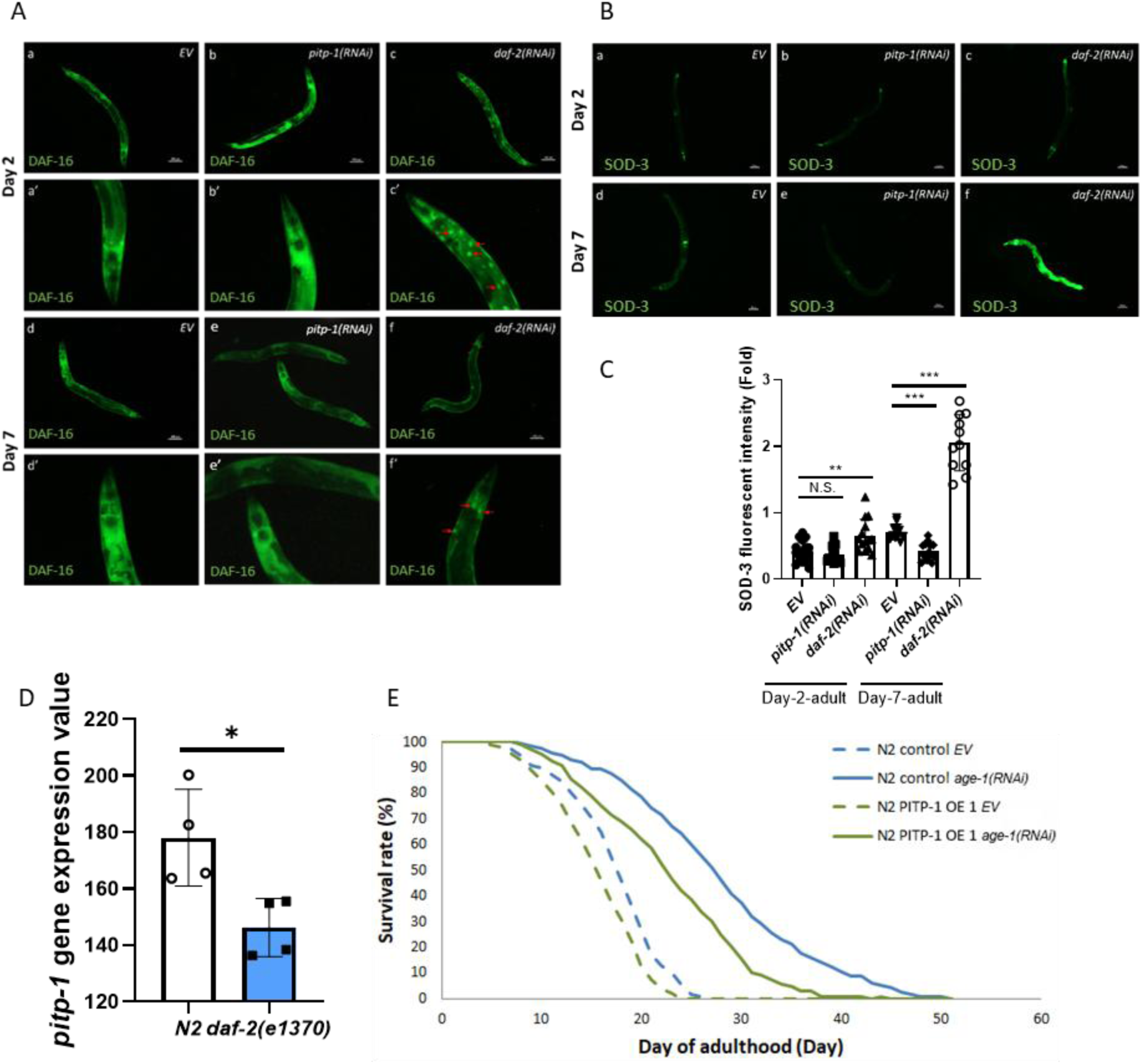
The role of *pitp-1* in IIS-mediated lifespan regulation. (A) Knockdown of *pitp-1* did not promote DAF-16 nuclear translocation. TJ356[*daf-16p::daf-16a/b::GFP + rol-6(su1006)*] was used as a DAF-16 reporter strain. Red arrows indicated DAF-16::GFP translocated into the nucleus and forms GFP puncta by *daf-2(RNAi)* as the positive control. (B, C) Knockdown of *pitp-1* did not increase *sod-3* expression. CF1553[*sod-3p::GFP + rol-6(su1006)*] was used as a *sod-3* reporter strain. (D) Whole-genome microarray data from *C. elegans* [30] revealed *pitp-1* expression was significantly reduced in *daf-2(e1370)*. (E) PITP-1 overexpression partially blocked the longevity effect by *age-1(RNAi)* knockdown. Survival curves are representative of three independent experiments. Statistical significance was determined by log-rank test for lifespan assays, ANOVA for multiple comparisons, and unpaired Student’s t-test where applicable.

## Supplementary Tables

**Table S1.**
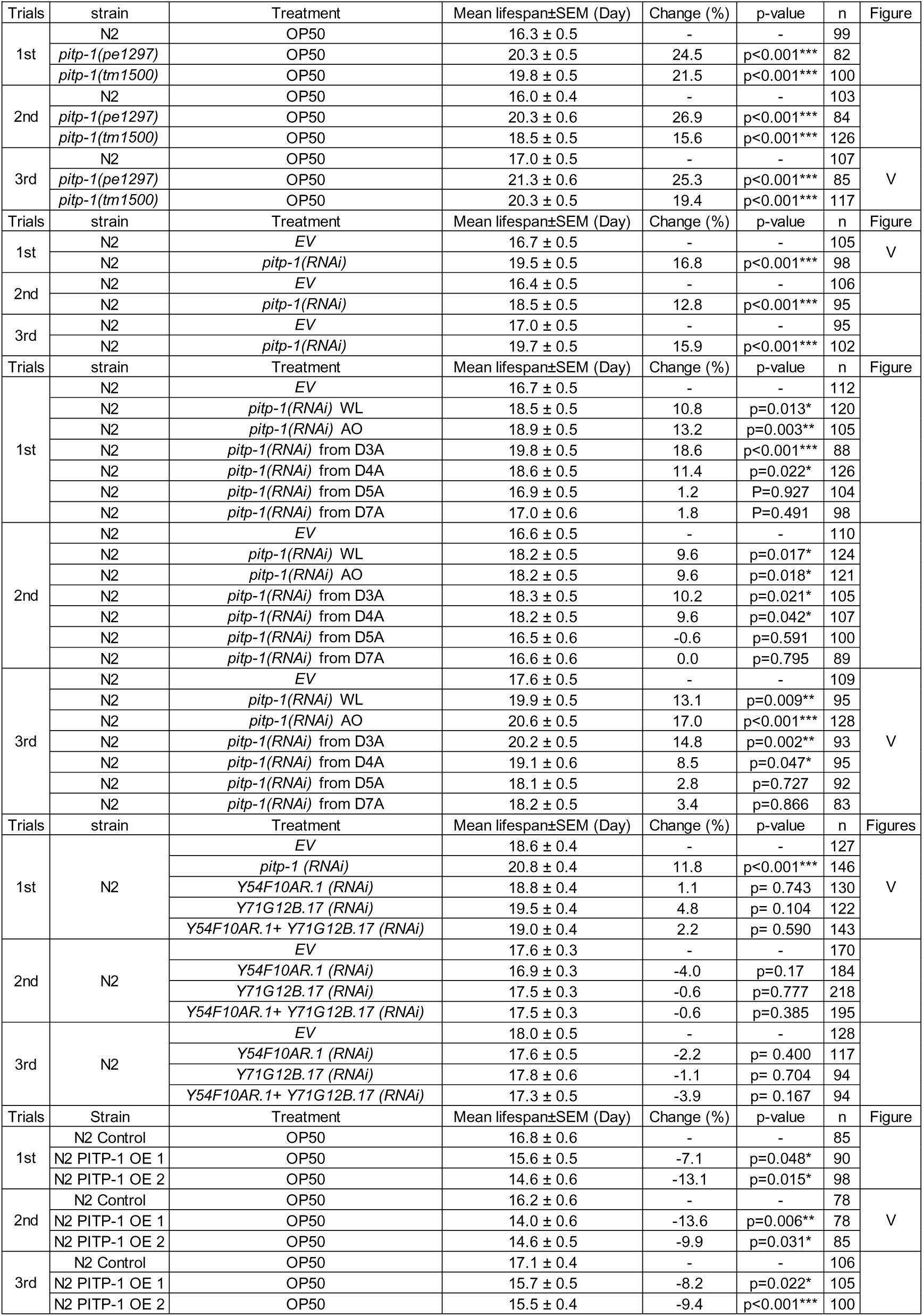
Lifespan analysis upon *pitp-1* inhibition and overexpression.

**Table S2.** Oxidative stress survival upon *pitp-1* inhibition.

| Table S2. Oxidative stress survival upon *pitp-1* inhibition. |  |  |  |  |  |  |  |  |
| --- | --- | --- | --- | --- | --- | --- | --- | --- |
| Trial | conc. | strain | Treatment | Mean lifespan ± SEM (Hours) | Compared with N2 | | n | Figure |
|  |  |  |  |  | Change (%) | p-value |  |  |
| 1st | 240uM Juglone | N2 | OP50 | 7.0 ± 0.4 | - | - | 87 | |
| | | *pitp-1(pe1297)* | OP50 | 9.3 ± 0.4 | 32.9 | p<0.001*** | 135 | |
| | | *pitp-1(tm1500)* | OP50 | 8.8 ± 0.4 | 25.7 | p=0.001** | 131 | |
| | 180uM Juglone | N2 | OP50 | 17.6 ± 1.0 | - | - | 90 | |
| | | *pitp-1(pe1297)* | OP50 | 23.7 ± 0.9 | 34.7 | p<0.001*** | 135 | |
| | | *pitp-1(tm1500)* | OP50 | 20.2 ± 1.0 | 14.8 | p=0.042* | 113 | |
| 2nd | 240uM Juglone | N2 | OP50 | 7.6 ± 0.3 | - | - | 140 | V |
| | | *pitp-1(pe1297)* | OP50 | 12.1 ± 0.6 | 59.2 | p<0.001*** | 128 | |
| | | *pitp-1(tm1500)* | OP50 | 10.4 ± 0.4 | 36.8 | p<0.001*** | 125 | |
| | 180uM Juglone | N2 | OP50 | 19.2 ± 0.9 | - | - | 130 | |
| | | *pitp-1(pe1297)* | OP50 | 29.9 ± 0.9 | 55.7 | p<0.001*** | 128 | |
| | | *pitp-1(tm1500)* | OP50 | 23.5 ± 1.0 | 22.4 | p<0.001*** | 121 | |
| Trial | conc. | strain | Treatment | Mean lifespan ± SEM (Hours) | Compared with EV | | n | Figure |
|  |  |  |  |  | Change (%) | p-value |  |  |
| 1st | 240uM Juglone | N2 | EV | 7.4 ± 0.3 | - | - | 108 | V |
| | | | *pitp-1(RNAi)* | 9.8 ± 0.4 | 32.4 | p<0.001*** | 109 | |
| | 180uM Juglone | N2 | EV | 17.5 ± 0.9 | - | - | 111 | |
| | | | *pitp-1(RNAi)* | 23.0 ± 1.0 | 31.4 | p<0.001*** | 109 | |
| 2nd | 240uM Juglone | N2 | EV | 8.8 ± 0.4 | - | - | 106 | |
| | | | *pitp-1(RNAi)* | 11.5 ± 0.5 | 30.7 | p<0.001*** | 105 | |
| | 180uM Juglone | N2 | EV | 18.1 ± 0.9 | - | - | 106 | |
| | | | *pitp-1(RNAi)* | 22.4 ± 1.0 | 23.8 | p<0.001*** | 105 | |

**Table S3.** Tissue-specific lifespan analysis upon *pitp-1* inhibition.

| Table S3. Tissue-specific lifespan analysis upon *pitp-1* inhibition. |  |  |  |  |  |  |  |
| --- | --- | --- | --- | --- | --- | --- | --- |
| Trials | Strain | Treatment | Mean lifespan±SEM (Day) | Change (%) | p-value | n | Figure |
| 1st | TU3401 | EV | 14.5 ± 0.4 | - | - | 109 | V |
|  |  | *pitp-1* (RNAi) | 17.4 ± 0.4 | 20 | p<0.001*** | 102 |  |
| 2nd | TU3401 | EV | 13.9 ± 0.3 | - | - | 101 |  |
|  |  | *pitp-1* (RNAi) | 16.2 ± 0.4 | 16.5 | p<0.001*** | 100 |  |
| 3rd | TU3401 | EV | 14.0 ± 0.3 | - | - | 105 |  |
|  |  | *pitp-1* (RNAi) | 16.5 ± 0.4 | 17.9 | p<0.001*** | 106 |  |
| Trials | Strain | Treatment | Mean lifespan±SEM (Day) | Change (%) | p-value | n | Figure |
| 1st | TU3311 | EV | 17.7 ± 0.5 | - | - | 107 | V |
|  |  | *pitp-1* (RNAi) | 20.8 ± 0.6 | 17.5 | p<0.001*** | 102 |  |
| 2nd | TU3311 | EV | 16.9 ± 0.4 | - | - | 104 |  |
|  |  | *pitp-1* (RNAi) | 19.5 ± 0.5 | 15.4 | p<0.001*** | 103 |  |
| 3rd | TU3311 | EV | 19.1 ± 0.5 | - | - | 109 |  |
|  |  | *pitp-1* (RNAi) | 21.2 ± 0.6 | 11 | p<0.001*** | 106 |  |
| Trials | Strain | Treatment | Mean lifespan±SEM (Day) | Change (%) | p-value | n | Figure |
| 1st | VP303 | EV | 14.5 ± 0.3 | - | - | 123 | V |
|  |  | *pitp-1* (RNAi) | 14.5 ± 0.3 | 0 | p=0.819 | 123 |  |
| 2nd | VP303 | EV | 14.0 ± 0.3 | - | - | 95 |  |
|  |  | *pitp-1* (RNAi) | 14.1 ± 0.3 | 0.7 | p=0.997 | 107 |  |
| 3rd | VP303 | EV | 14.0 ± 0.3 | - | - | 111 |  |
|  |  | *pitp-1* (RNAi) | 14.1 ± 0.4 | 0.7 | p=0.593 | 109 |  |
| Trials | Strain | Treatment | Mean lifespan±SEM (Day) | Change (%) | p-value | n | Figure |
| 1st | WM118 | EV | 15.5 ± 0.5 | - | - | 109 | V |
|  |  | *pitp-1* (RNAi) | 16.0 ± 0.5 | 3.2 | p=0.345 | 120 |  |
| 2nd | WM118 | EV | 15.4 ± 0.4 | - | - | 111 |  |
|  |  | *pitp-1* (RNAi) | 15.7 ± 0.5 | 2.6 | p=0.195 | 106 |  |
| 3rd | WM118 | EV | 15.9 ± 0.6 | - | - | 92 |  |
|  |  | *pitp-1* (RNAi) | 16.3 ± 0.5 | 2.5 | p=0.827 | 93 |  |
| Trials | Strain | Treatment | Mean lifespan±SEM (Day) | Change (%) | p-value | n | Figure |
| 1st | XE1375 | EV | 13.9 ± 0.2 | - | - | 127 |  |
|  |  | *pitp-1* (RNAi) | 15.1 ± 0.2 | 8.6 | p= 0.004** | 146 |  |
| 2nd | XE1375 | EV | 14.8 ± 0.2 | - | - | 119 | V |
|  |  | *pitp-1* (RNAi) | 16.2 ± 0.3 | 9.5 | p= 0.004** | 118 |  |
| 3rd | XE1375 | EV | 14.0 ± 0.3 | - | - | 135 |  |
|  |  | *pitp-1* (RNAi) | 15.2 ± 0.1 | 8.6 | p= 0.029* | 118 |  |
| Trials | Strain | Treatment | Mean lifespan±SEM (Day) | Change (%) | p-value | n | Figure |
| 1st | XE1582 | EV | 12.5 ± 0.3 | - | - | 112 | V |
|  |  | *pitp-1* (RNAi) | 13.8 ± 0.6 | 10.4 | p=0.002** | 110 |  |
| 2nd | XE1582 | EV | 13.2 ± 0.1 | - | - | 106 |  |
|  |  | *pitp-1* (RNAi) | 14.4 ± 0.4 | 9.1 | p=0.030* | 102 |  |
| 3rd | XE1582 | EV | 13.3 ± 0.4 | - | - | 110 |  |
|  |  | *pitp-1* (RNAi) | 14.7 ± 0.3 | 10.5 | p=0.003** | 104 |  |
| Trials | Strain | Treatment | Mean lifespan±SEM (Day) | Change (%) | p-value | n | Figure |
| 1st | XE1581 | EV | 11.8 ± 0.6 | - | - | 106 | V |
|  |  | *pitp-1* (RNAi) | 12.7 ± 0.3 | 7.6 | p= 0.004** | 114 |  |
| 2nd | XE1581 | EV | 12.4 ± 0.3 | - | - | 119 |  |
|  |  | *pitp-1* (RNAi) | 13.2 ± 0.4 | 6.5 | p= 0.011* | 127 |  |
| 3rd | XE1581 | EV | 14.0 ± 0.4 | - | - | 186 |  |
|  |  | *pitp-1* (RNAi) | 15.0 ± 0.3 | 7.1 | p= 0.012* | 155 |  |
| Trials | Strain | Treatment | Mean lifespan±SEM (Day) | Change (%) | p-value | n | Figure |
| 1st | XE1474 | EV | 12.9 ± 0.3 | - | - | 108 |  |
|  |  | *pitp-1* (RNAi) | 13.6 ± 0.3 | 5.4 | p=0.239 | 111 |  |
| 2nd | XE1474 | EV | 13.5 ± 0.3 | - | - | 106 | V |
|  |  | *pitp-1* (RNAi) | 14.0 ± 0.3 | 3.7 | p=0.097 | 105 |  |

**Table S4.**
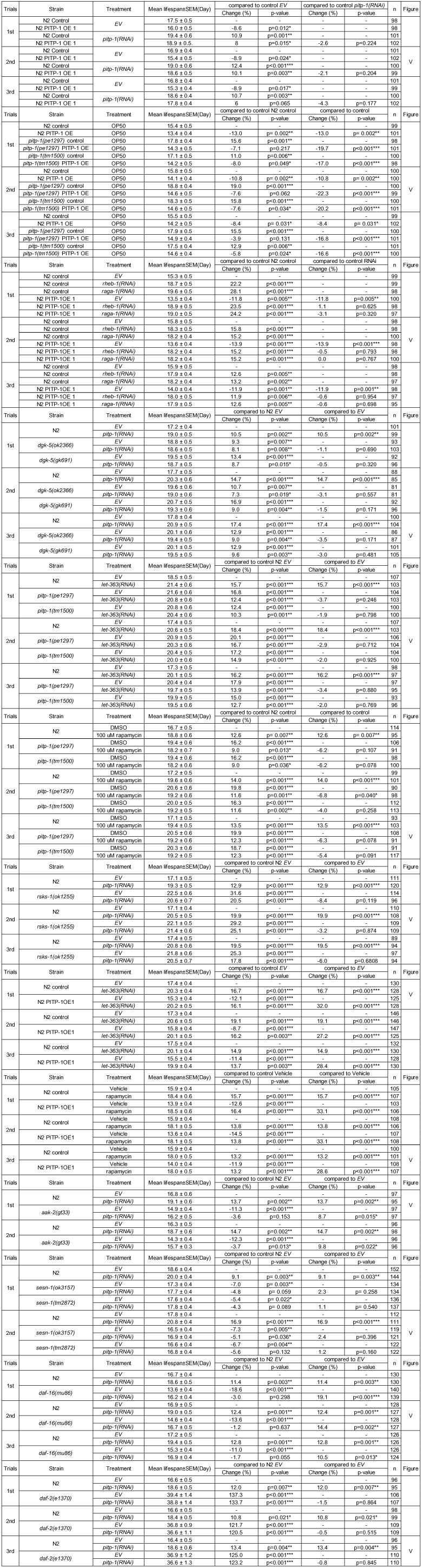
Genetic epistasis tests of *pitp-1* regulation on lifespan.

**Table S5.**
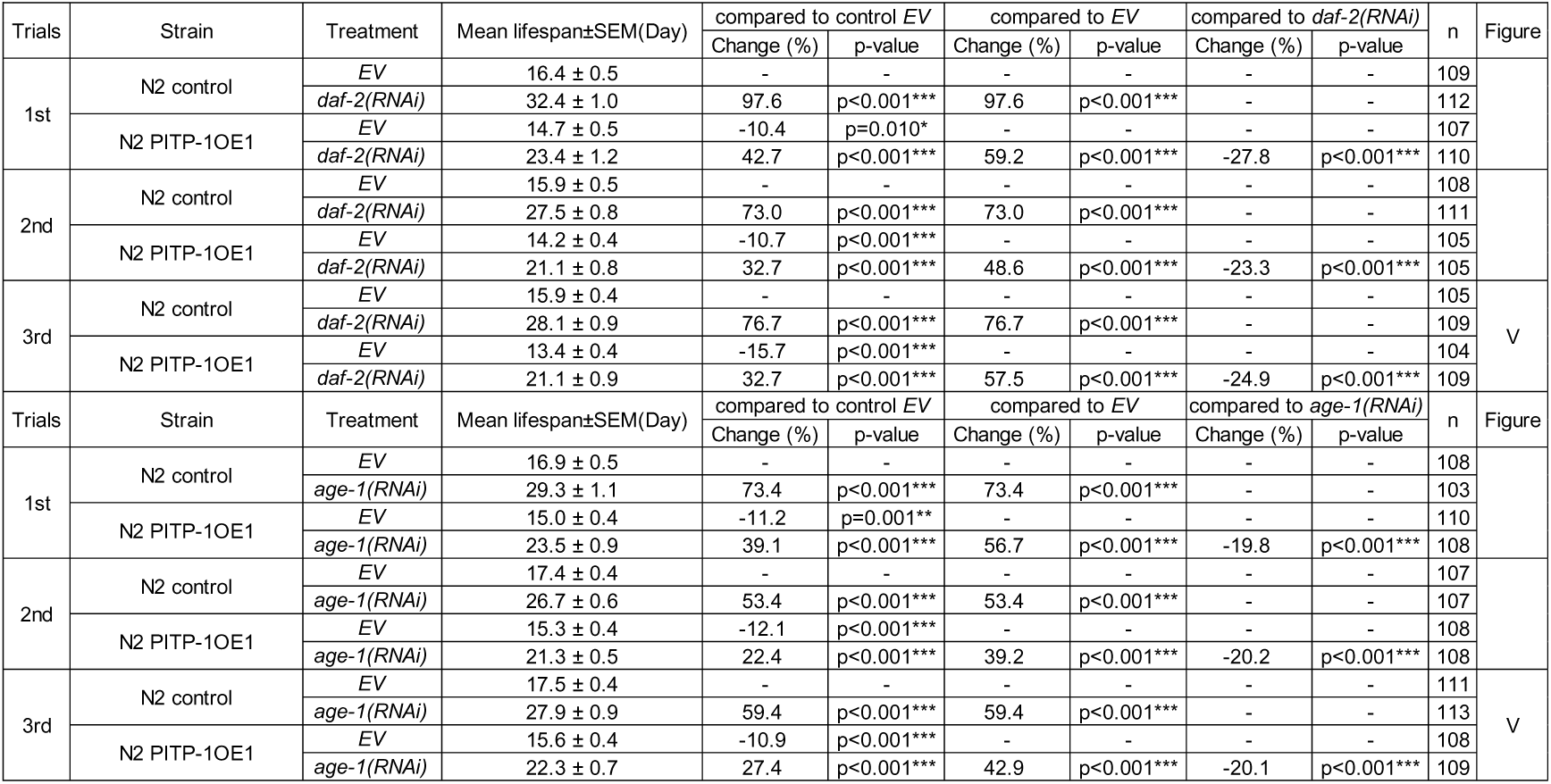
Lifespan epistasis analysis of *pitp-1* overexpression with IIS inhibition.

|  |  |  |  |  |  |  |  |  |  |  |  |
| --- | --- | --- | --- | --- | --- | --- | --- | --- | --- | --- | --- |
| Trials | Strain | Treatment | Mean lifespan±SEM(Day) | compared to control *EV* | | compared to *EV* | | compared to *daf-2(RNAi)* | | n | Figure |
|  |  |  |  | Change (%) | p-value | Change (%) | p-value | Change (%) | p-value |  |  |
| 1st | N2 control | *EV* | 16.4 ± 0.5 | - | - | - | - | - | - | 109 | |
| | | *daf-2(RNAi)* | 32.4 ± 1.0 | 97.6 | p<0.001*** | 97.6 | p<0.001*** | - | - | 112 | |
| | N2 PITP-1OE1 | *EV* | 14.7 ± 0.5 | -10.4 | p=0.010* | - | - | - | - | 107 | |
| | | *daf-2(RNAi)* | 23.4 ± 1.2 | 42.7 | p<0.001*** | 59.2 | p<0.001*** | -27.8 | p<0.001*** | 110 | |
| 2nd | N2 control | *EV* | 15.9 ± 0.5 | - | - | - | - | - | - | 108 | |
| | | *daf-2(RNAi)* | 27.5 ± 0.8 | 73.0 | p<0.001*** | 73.0 | p<0.001*** | - | - | 111 | |
| | N2 PITP-1OE1 | *EV* | 14.2 ± 0.4 | -10.7 | p<0.001*** | - | - | - | - | 105 | |
| | | *daf-2(RNAi)* | 21.1 ± 0.8 | 32.7 | p<0.001*** | 48.6 | p<0.001*** | -23.3 | p<0.001*** | 105 | |
| 3rd | N2 control | *EV* | 15.9 ± 0.4 | - | - | - | - | - | - | 105 | V |
| | | *daf-2(RNAi)* | 28.1 ± 0.9 | 76.7 | p<0.001*** | 76.7 | p<0.001*** | - | - | 109 | |
| | N2 PITP-1OE1 | *EV* | 13.4 ± 0.4 | -15.7 | p<0.001*** | - | - | - | - | 104 | |
| | | *daf-2(RNAi)* | 21.1 ± 0.9 | 32.7 | p<0.001*** | 57.5 | p<0.001*** | -24.9 | p<0.001*** | 109 | |
| Trials | Strain | Treatment | Mean lifespan±SEM(Day) | compared to control *EV* | | compared to *EV* | | compared to *age-1(RNAi)* | | n | Figure |
|  |  |  |  | Change (%) | p-value | Change (%) | p-value | Change (%) | p-value |  |  |
| 1st | N2 control | *EV* | 16.9 ± 0.5 | - | - | - | - | - | - | 108 | |
| | | *age-1(RNAi)* | 29.3 ± 1.1 | 73.4 | p<0.001*** | 73.4 | p<0.001*** | - | - | 103 | |
| | N2 PITP-1OE1 | *EV* | 15.0 ± 0.4 | -11.2 | p=0.001** | - | - | - | - | 110 | |
| | | *age-1(RNAi)* | 23.5 ± 0.9 | 39.1 | p<0.001*** | 56.7 | p<0.001*** | -19.8 | p<0.001*** | 108 | |
| 2nd | N2 control | *EV* | 17.4 ± 0.4 | - | - | - | - | - | - | 107 | |
| | | *age-1(RNAi)* | 26.7 ± 0.6 | 53.4 | p<0.001*** | 53.4 | p<0.001*** | - | - | 107 | |
| | N2 PITP-1OE1 | *EV* | 15.3 ± 0.4 | -12.1 | p<0.001*** | - | - | - | - | 108 | |
| | | *age-1(RNAi)* | 21.3 ± 0.5 | 22.4 | p<0.001*** | 39.2 | p<0.001*** | -20.2 | p<0.001*** | 108 | |
| 3rd | N2 control | *EV* | 17.5 ± 0.4 | - | - | - | - | - | - | 111 | V |
| | | *age-1(RNAi)* | 27.9 ± 0.9 | 59.4 | p<0.001*** | 59.4 | p<0.001*** | - | - | 113 | |
| | N2 PITP-1OE1 | *EV* | 15.6 ± 0.4 | -10.9 | p<0.001*** | - | - | - | - | 108 | |
| | | *age-1(RNAi)* | 22.3 ± 0.7 | 27.4 | p<0.001*** | 42.9 | p<0.001*** | -20.1 | p<0.001*** | 109 | |

## Acknowledgments

We thank the *C. elegans* Core Facility of the National Core Facility for biopharmaceuticals, National Science and Technology Council (NSTC), in Taiwan for technical support, and the assistance from Dr. Ao-Lin Hsu. We thank the technical support from Ya-Hsien Chou at the confocal imaging core at National Tsing Hua University sponsored by NSTC.

## Author contribution

H. D. Wang, C. H. Yuh, and Y. H. Lin contributed to the conception, design of the study. Y. H. Lin and H. D. Wang wrote the manuscript. H. D. Wang and C. H. Yuh contributed to funding acquisition. Y. H. Lin, Y. H. Liao, S. B. Liao, T. Y. Lin and P. J. Hsu contributed to the acquisition of data and helped the data analysis. Y. H. Lin, H. D. Wang, C. S. Chen, T. T. Ching and C. H. Yuh contributed to the development of methodology. C. H. Yuh, Y. H. Lin and H. D. Wang contributed to analyze transcriptomic profiles. C. S. Chen, T. T. Ching and O. I. Wagner offered RNAi clones or *C. elegans* strains. Y. H. Lin, T. T. Ching, M. M. Shanmugam and O. I. Wagner contributed to establish overexpression construct and microinject transgenic strains. Y. H. Lin, H. D. Wang, C. S. Chen and C. H. Yuh contributed to the interpretation of data.

## Corresponding authors

Correspondence to Horng-Dar Wang and Chiou-Hwa Yuh

## Ethics declarations

### Ethics approval and consent to participate

Not applicable. The analysis of *PITPNM1, PITPNM2,* and *PITPNM3* expressions was from the RNAseq raw data in the published GEO, no human tissue was used.

### Consent for publication

Not applicable.

